# Engineering SARS-CoV-2 neutralizing antibodies for increased potency and reduced viral escape

**DOI:** 10.1101/2022.01.06.475303

**Authors:** Fangzhu Zhao, Celina Keating, Gabriel Ozorowski, Namir Shaabani, Irene M. Francino-Urdaniz, Shawn Barman, Oliver Limbo, Alison Burns, Panpan Zhou, Michael J. Ricciardi, Jordan Woehl, Quoc Tran, Hannah L. Turner, Linghang Peng, Deli Huang, David Nemazee, Raiees Andrabi, Devin Sok, John R. Teijaro, Timothy A. Whitehead, Andrew B. Ward, Dennis R. Burton, Joseph G. Jardine

## Abstract

The rapid spread of SARS-CoV-2 variants poses a constant threat of escape from monoclonal antibody and vaccine countermeasures. Mutations in the ACE2 receptor binding site on the surface S protein have been shown to disrupt antibody binding and prevent viral neutralization. Here, we use a directed evolution-based approach to engineer three neutralizing antibodies for enhanced binding to S protein. The engineered antibodies showed increased *in vitro* functional activity in terms of neutralization potency and/or breadth of neutralization against viral variants. Deep mutational scanning revealed that higher binding affinity reduced the total number of viral escape mutations. Studies in the Syrian hamster model showed two examples where the affinity matured antibody provided superior protection compared to the parental antibody. These data suggest that monoclonal antibodies for anti-viral indications could benefit from *in vitro* affinity maturation to reduce viral escape pathways and appropriate affinity maturation in vaccine immunization could help resist viral variation.

## INTRODUCTION

Over the last year and a half, severe acute respiratory syndrome coronavirus 2 (SARS-CoV-2) has had devastating consequences for global health and economies. Following the discovery of the disease, there was a rush to produce protective vaccines and therapeutics. Multiple highly effective vaccines have been developed that elicit immune responses against the SARS-CoV-2 spike (S) trimer^1^. The protective mechanisms for the coronavirus disease 2019 (COVID-19) vaccines are still being deduced, however, several analyses have found that the elicitation of neutralizing antibodies (nAbs) correlates with protection^2, 3^, a finding consistent with many other successful antiviral vaccines^4^. NAbs have been identified that target several distinct epitopes on the S trimer, but the majority of nAbs target the receptor binding domain (RBD)^5, 6^. While vaccines are undisputedly the most effective strategy for control of COVID-19, recombinantly produced nAbs offer the potential to supplement prophylactic coverage in populations that respond poorly to vaccines, e.g. immunocompromised individuals, can be administered as a post-exposure prophylactic and can be used therapeutically to prevent hospitalization^7, 8^.

One of the unique challenges in using a neutralizing monoclonal antibody (mAb) for antiviral indications is addressing existing viral diversity and the high mutational propensity in viruses that can give rise to resistant viral variants. Since the discovery of SARS-CoV-2 in 2019, thousands of viral variants containing synonymous and nonsynonymous mutations have been documented^9^. A growing number of these new variants (termed “Variants of Concern” or VOCs)^10^ contain mutations that increase infectivity and/or allow viral escape from monoclonal nAbs elicited against the original SARS-CoV-2^11–14^. Several strategies are commonly used to mitigate the likelihood of viral escape from nAbs. Investigators often select antibodies that target functionally important and conserved regions, reducing the number of mutations that can allow viral escape without incurring a fitness cost^15–17^. It is also common to use cocktails of at least two nAbs targeting different epitopes, so multiple mutations are necessary for viral escape^18–21^. A third approach that is less well explored is to *in vitro* affinity mature the nAb against the target antigen to increase the binding affinity, helping to mitigate the impact of the viral mutations^22^. Here, we explore how increased binding affinity impacts the *in vitro* neutralization breadth and potency of three COVID-19 nAbs, CC12.1, CC6.30 and CC6.33^23^, and how these affinity improvements impact the *in vivo* protective capability of these nAbs.

## RESULTS

### Structural analysis of nAbs CC6.30 and CC6.33

Previously, we reported the structure of nAb CC12.1, which binds to the RBS-A or Class 1 epitope site and competes directly with angiotensin-converting enzyme 2 (ACE2)^5, 24, 25^ but specificities of two other nAbs of interest, CC6.30 and CC6.33, were limited to epitope-binning data^23^. To better understand the molecular contacts of antibodies, we used cryoelectron microscopy (cryoEM) to solve the structures of two of the antibodies in complex with stabilized SARS-CoV-2 S trimers: 1) nAb CC6.30, which targets the RBS-B or Class 2 epitope site, binds RBD with an affinity of 1.7 nM (IgG format) and directly competes with ACE2, and 2) nAb CC6.33, which targets the Class 3 epitope site that is distinct from the ACE2 binding site and binds RBD with an affinity of 257 nM (Fig.1 and Fig.2). Several clinical-stage nAbs also recognize these epitopes such as CB6/LY- CoV16 (RBS-A or Class 1)^25, 26^, REGN10933 (RBS-B or Class 2)^14, 27^ and S309 (Class 3)^5, 28^. In each complex, we used a stabilized, uncleaved Spike trimer (HPM7) based on the Wuhan strain containing 6 proline (hexaproline or HP) mutations^29^ and an engineered interprotomer disulfide (mut7 or M7) between residues 705 and 883 of the S2 subunit. The global resolution of the two complexes was 3.6 Å (CC6.30/HPM7) and 3.3 Å (CC6.33/HPM7) and model building was assisted by local refinement maps of just the RBD and fragment antigen binding (Fab) variable region (Supplementary Fig.1,2, and Supplementary Table 1,2). Additionally, we solved the ligand-free structure of the HPM7 spike (2.8 Å and 3.0 Å resolution for C3 symmetric and asymmetric, respectively) (Supplementary Fig.2c,d and Supplementary Table 2).

**Fig. 1.**
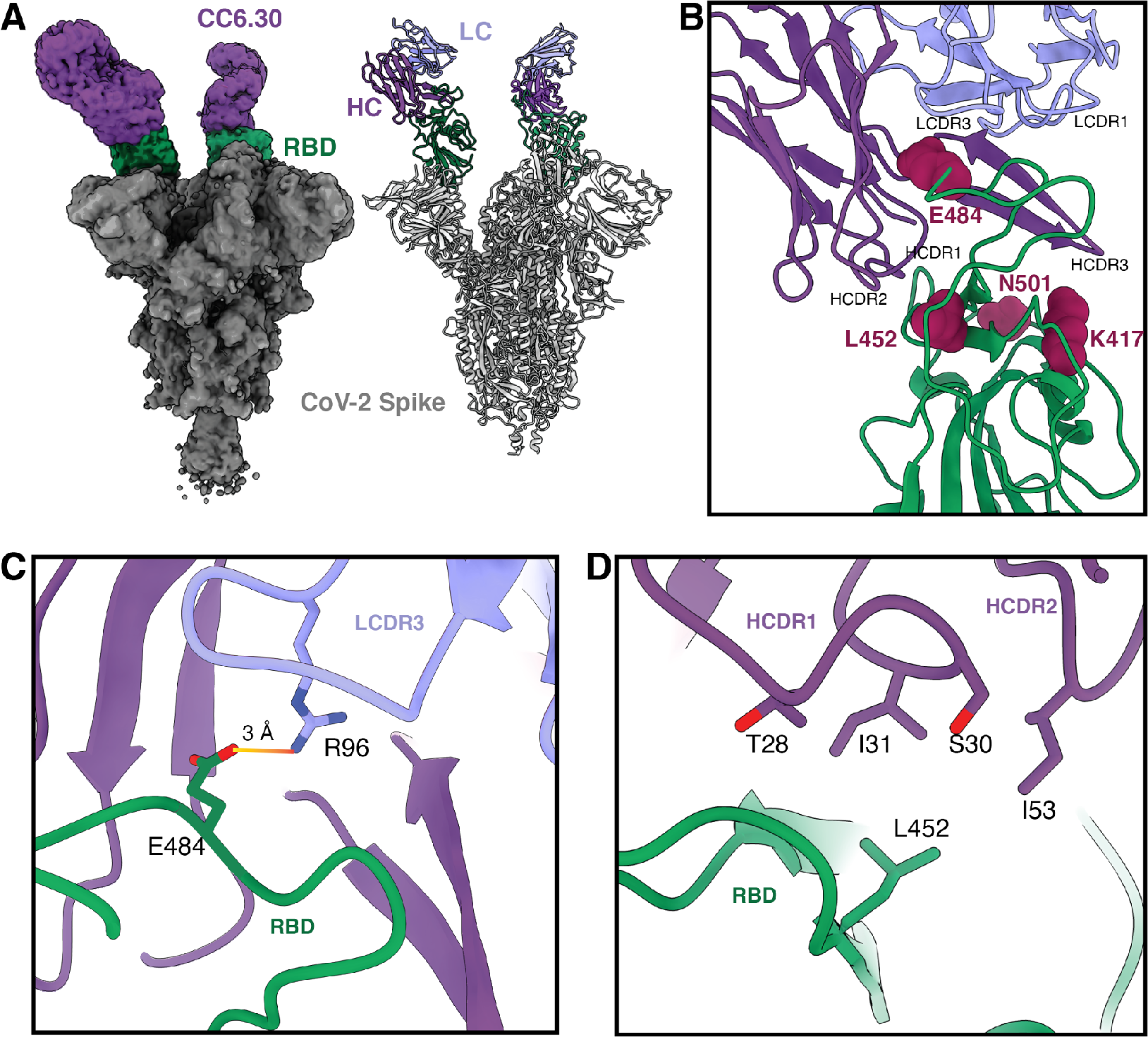
Structural characterization of nAb CC6.30. **a**, CryoEM map and model, with CC6.30 Fab and Spike RBD colored for clarity. **b**, Overview of CC6.30/RBD interface with key antibody CDRs and SARS-CoV-2 VOC mutation sites labeled. **c**, Salt bridge between LCDR3 R96 and RBD E484. **d**, Hydrophobic interactions between RBD L452 and side chains of HCDR1 and HCDR2.

**Fig. 2.**
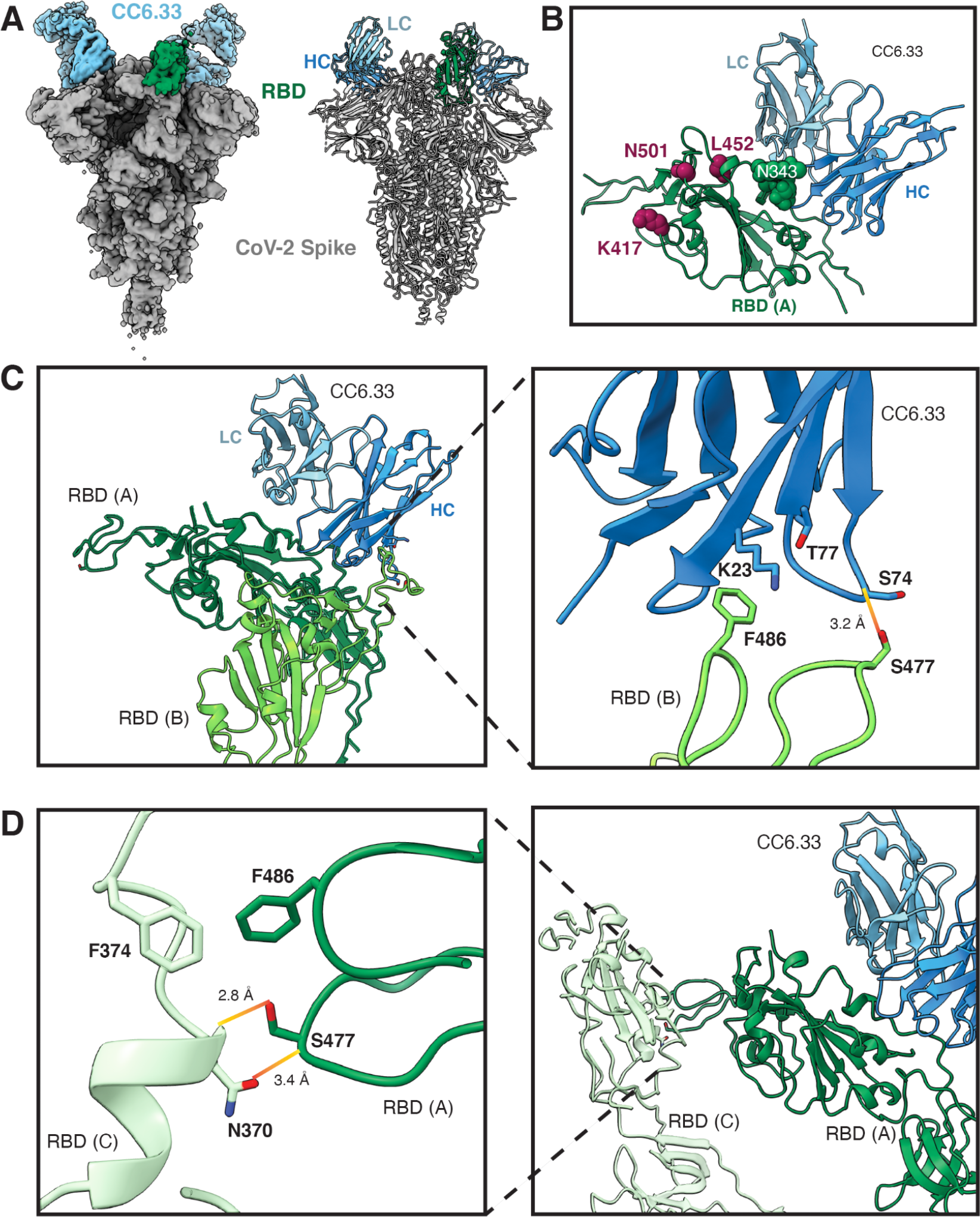
Structural characterization of nAb CC6.33. **a**, CryoEM map and model, with CC6.33 Fab and Spike RBD colored for clarity. **b**, Overview of CC6.33/RBD interface that is centered on the Spike N343 glycan. Common mutation sites in VOCs are highlighted and are distal from the CC6.33 epitope. The RBD ridge of a neighboring protomer interacts with framework regions of CC6.33 **c** in a manner similar to the ridge of a down-RBD supporting a neighboring up-RBD **d**.

CC6.30 straddles the SARS-CoV-2 RBD ridge (approximately residues 473-490) with contributions from all heavy chain (HC) and light chain (LC) complementarity determining regions (CDRs) except for LCDR2 (Fig.1a,b). The HCDR3 contains a disulfide bond (C99-C100d, Kabat numbering) that makes it more rigid, perhaps in turn acting to stabilize this more flexible region of the RBD around the “ACE2-binding ridge” (approximately Spike residue numbers 470-490) (Supplementary Fig.1b,c). Important interactions at the binding interface include hydrophobic packing of the RBD F490 side chain with antibody HC side chains I31, I52, I53 and I54 (Supplementary Fig.1d), and predicted hydrogen bonding between the side chain of HCDR3 R97 and both the RBD I492 backbone carbonyl and Q493 side chain (Supplementary Fig.1e). A salt bridge is formed between the side chains of RBD E484 and LCDR3 R96 (Fig.1c). This interaction is predicted to decrease the effectiveness of CC6.30 against certain VOCs, specifically Beta^30^ and Gamma^31^ VOCs, which contain an E484K mutation that abolishes the salt bridge and causes a possible charge-charge repulsion. Of similar concern, the L452 side chain of the RBD has hydrophobic interactions with HCDR1 I31 and HCDR2 I53 which would be incompatible with a long, positively-charged side chain that results from the L452R mutation found in the Delta variant^32^ (Fig.1d).

CC6.30 appears to only bind the RBD-up state of the Spike (Fig.1a). While the ligand-free cryoEM structures of HPM7 spike reveal a 3 RBD-down conformation, our data suggest that the antibody is capable of shifting the equilibrium by capturing RBD in the up conformation. The most stable cryoEM reconstruction contains 2 up RBDs, each bound by CC6.30, while the unoccupied RBD remains in the down position (Fig.1a). Superposition of the RBD:CC6.30 portion of the model onto the all-down RBD ligand-free structure predicts that clashes would occur between HCDR3 and the RBD and N343 glycan from an adjacent protomer (Supplementary Fig.1f). Finally, E484 is a major part of the epitope as defined in the Class 2 or RBS-B nomenclatures^14^, consistent with our other observations for CC6.30.

CC6.33 binds a non-overlapping epitope to that of CC6.30, with the heavy and light chain interface centered on the N343 glycan (Supplementary Fig.2a,b). In contrast to CC6.30, this nAb binds the RBD-down conformation and the most stable reconstruction has 2 down RBDs bound by the antibody, while the third RBD is in the up position (Fig.2a). Indeed, the binding of CC6.33 to a down RBD requires slight opening of the apex to relieve a clash with the RBD ridge of the adjacent protomer, and is likely the driving force for the unoccupied RBD shifting to the up position after the binding of two Fabs (Supplementary Fig.2e). While modeling suggests that CC6.33 should be able to bind an up RBD, we did not observe this in our dataset, possibly due to the HPM7 spike design preferentially displaying 3 down-RBDs (Supplementary Fig.2f). Also, portions of HC framework regions 1 (HFR1) and HFR3 contact the RBD ridge of the neighboring protomer (still in the down position) in a manner that mimics the interaction between the unoccupied up-RBD and an adjacent RBD-down ridge, further stabilizing the interaction between antibody and spike (Fig.2c,d). Binding of CC6.33 to its epitope is largely governed by hydrophobic interactions involving the HC, including HCDR2 residues I52, I53 and L54 packing against RBD residues L335, V362 and P527 (Supplementary Fig.2g). HCDR3 W98 reaches into an aromatic pocket lined with RBD residues F338, F342, A363, Y365 and L368, while also donating a hydrogen bond to the backbone carbonyl of D364 (Supplementary Fig.2h). Fewer hydrogen bonds are predicted between RBD and CC6.33 compared to CC6.30. Those contributed by CC6.33 often involve bonds to RBD main chain atoms (e.g. HCDR3 Q97 with RBD backbone C336, V362 and D364), making such interactions less susceptible to changes in side chains resulting from VOC mutations (Supplementary Fig.2i). The antibody epitope itself is largely positioned away from the common RBD mutations that could affect binding, a property shared with other Class 3 RBD antibodies (Fig.2d). Lastly, the LC is mostly involved via LCDR2 packing against and providing hydrogen bonds to the viral N343 glycan, and a single peptide-peptide hydrogen bond between the side chains of LCDR1 Y32 and RBD E340 (Fig.2b and Supplementary Fig.2j).

### Engineering higher affinity SARS-CoV-2 antibodies

Affinity maturation of CC12.1, CC6.30 and CC6.33 was achieved using our rapid affinity maturation strategy, SAMPLER^33^. Briefly, HC and LC libraries were synthesized containing one mutation per CDR loop from the starting sequence, for up to three mutations per chain. Potential liabilities were informatically filtered from the library process and an N-linked glycan on LCDR1 of CC6.30 was removed by an N28S mutation, reverting that position to the original amino acid found in the germline VK1-39 gene segment so that any improved CC6.30 variant would not contain that glycan. The HC and LC libraries were displayed on the surface of yeast and iterative rounds of selections were used to enrich for clones with higher affinity for SARS-CoV-2 RBD (for CC12.1 and CC6.30 libraries) or S protein (for the CC6.33 library). The sort process also included a round of negative selection, where clones with low binding to a polyclonal preparation of detergent-solubilized HEK293 cell membrane proteins were enriched to remove polyreactive variants. The enriched clones were then combined into a heavy/light combinatorial library and screened again with the same four-round selection strategy to identify the optimal heavy/light pairs^33^. At the conclusion of the selection process, sequences of the antibodies were recovered and 12 improved variants of each antibody were selected to be reformatted and expressed as IgG for characterization.

All enhanced (e) eCC12.1, eCC6.30 and eCC6.33 variants bound SARS-CoV-2 RBD with higher affinity than the parental antibodies, with an average 45-fold (5- to 267-fold) increase in monovalent equilibrium dissociation constants (Fig.3a) measured by surface plasmon resonance (SPR). In nearly all cases, the affinity gains came through a reduction in the dissociation rate (Supplementary Table 3). The binding affinity for monomeric RBD is notably lower for CC6.33 compared to CC12.1, CC6.30 (Fig.3a) and the majority of other antibodies isolated from our COVID- 19 cohort^23^. The binding affinity of CC6.33 Fab for S protein is approximately 10-fold higher than for RBD, suggesting that the CC6.33 epitope is poorly formed on monomeric RBD and/or differential processing of the N343 glycan affects mAb binding. ELISA binding to SARS-CoV-2 RBD and S by CC12.1 and CC6.30 parental and engineered nAbs (enAbs) was comparable, however, a large improvement in neutralization EC50 and the maximum neutralization plateau was observed for eCC6.33 variants compared to the CC6.33 parental (Supplementary Fig.3). The enAb variants were evaluated by analytical size exclusion chromatography and found to be monodispersed with similar column retention time to our clinical controls (Supplementary Fig.4). None of the eCC12.1 or eCC6.33 variants bound to antigens in our polyreactivity panel (Chinese hamster ovary cell solubilized membrane proteins, single-stranded DNA, and human insulin) or stained HEp2 epithelial cells (Supplementary Fig.5). Several of the CC6.30 variants showed low levels of binding to one or more of the antigens in our polyreactivity panel or stained HEp2 cells, but the majority of engineered variants were negative in all assays, highlighting the importance of expressing and validating multiple variants.

**Fig. 3.**
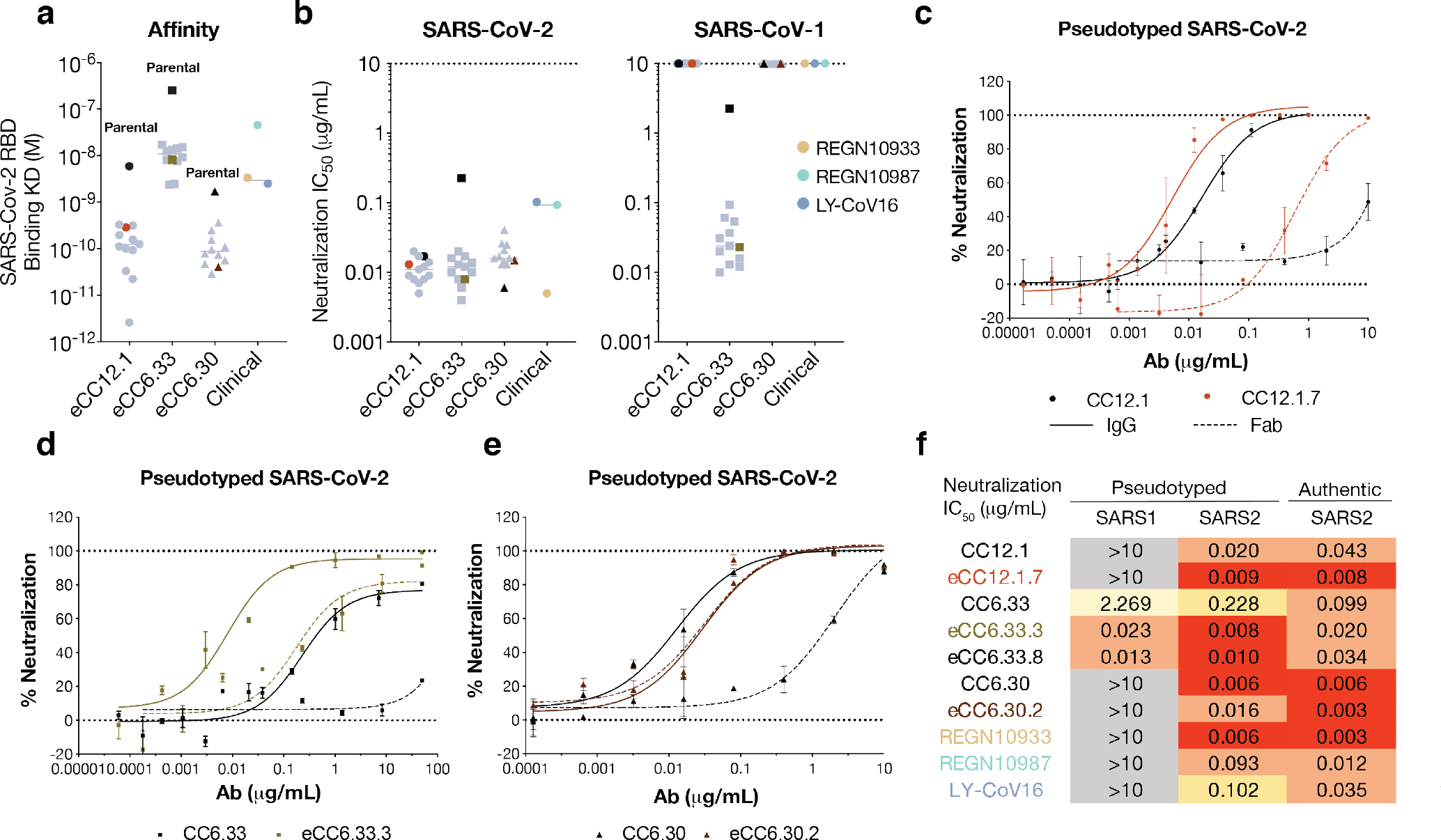
Binding affinity and neutralization potency of engineered SARS-CoV-2 nAbs. **a**, Enhanced and parental nAbs binding affinity against SARS-CoV-2 RBD by surface plasmon resonance. Parental nAbs are highlighted in black. eCC12.1.7, eCC6.33.3 and eCC6.30.2 are highlighted whereas other engineered variants are colored in grey. RBD binding to antibodies via a Fc-capture, multi-cycle method. Association and dissociation rate constants were calculated through a 1:1 Langmuir binding model using the BIAevaluation software. **b**, Neutralization IC50 against pseudotyped SARS-CoV-2 and SARS-CoV-1 viruses. **c-e**, SARS-CoV-2 pseudovirus neutralization curves of **c** parental CC12.1 and eCC12.1.7, **d** parental CC6.33 and eCC6.33.3, **e** parental CC6.30 and eCC6.30.2 in IgG and Fab molecules. Solid lines represent IgG neutralization whereas dashed lines represent Fab neutralization. Assays were run in duplicate. Error bars represent standard deviation. Data are representative of at least two independent experiments. **f**, Summary table of nAb neutralization IC50 against pseudotyped SARS-CoV-1 and SARS-CoV2, as well as replicating SARS-CoV-2.

**Fig. 4.**
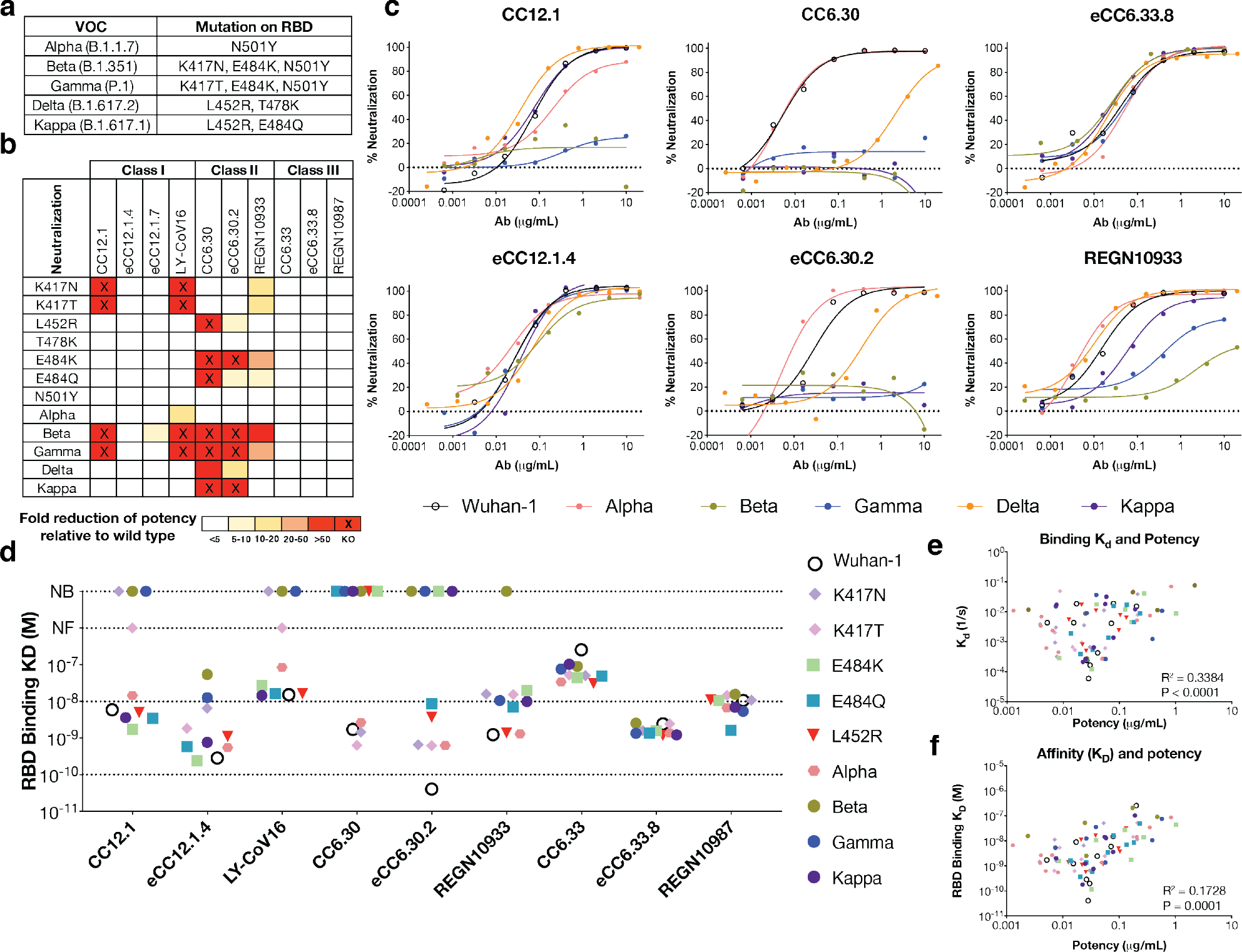
Antibody neutralization activities against circulating variants. **a**, Circulating SARS-CoV- 2 variants of concern (VOC) and their mutations on RBD. **b**, Summary table of fold reduction of neutralization potency of mAbs against circulating SARS-CoV-2 variants and single mutations relative to the original lineage Wuhan-1. Fold difference of neutralization potency was colored according to the key. **c**, Representative neutralization curves of CC12.1, eCC12.1.7, CC6.30, eCC6.30.2, eCC6.33.8 and REGN10933 against Wuhan-1 as well as VOC including Alpha (B.1.1.7), Beta (B.1.351), Gamma (P.1), Delta (B.1.617.2), Kappa (B.1.617.1). Assays were run in duplicate. Data are representative of at least two independent experiments. SARS-CoV-2 and variants were colored according to the key. **d**, Antibody binding affinity (KD) against wild type SARS- CoV-2 RBD and RBD mutant proteins. NB: no binding. NF: Not fit to a simple kinetics model. Antibodies were captured to SPR sensors via a Fc-capture, multi-cycle method. Association and dissociation rate constants were calculated through a 1:1 Langmuir binding model using the BIAevaluation software. **e-f,** Pearson correlation analysis between antibody **e** off-rate constant (Kd) or **f** equilibrium dissociation constant (KD) binding kinetics against RBD variants and antibody neutralization potency against mutant pseudoviruses.

**Fig. 5.**
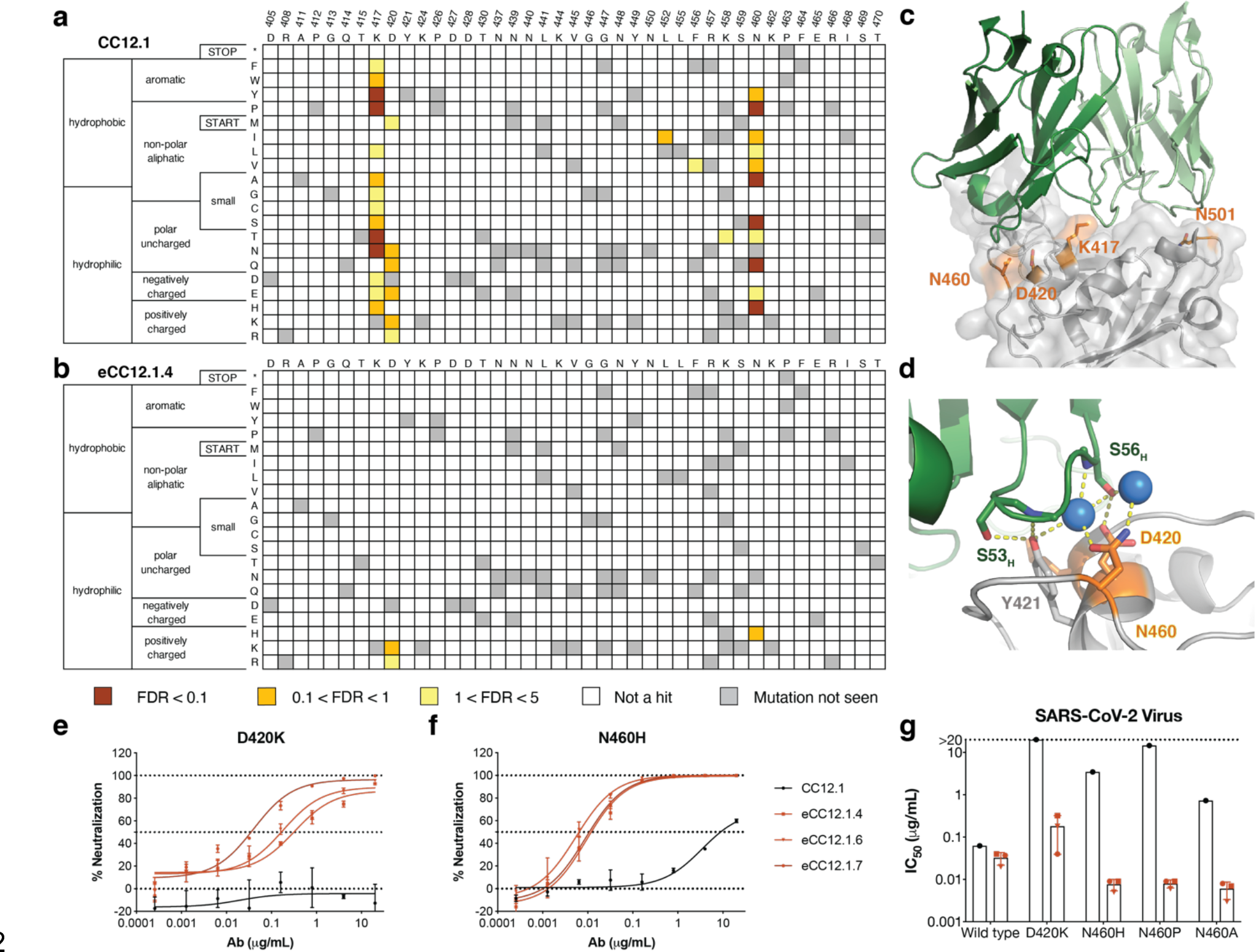
Mapping potential RBD viral escape mutants from CC12.1 and eCC12.1.4. **a-b**, Putative viral escape screening of **a** CC12.1 and **b** eCC12.1.4 using a RBD yeast display platform ^36^. RBD variants that did not disrupt ACE2 interaction but evaded nAb recognition were sorted and sequenced. A control with no ACE2 labeling was also sorted and served as an empirical false discovery rate (FDR). The enrichment ratio for each mutation relative to the reference population was colored according to the key. The heatmap of RBD residues 405 - 470 was shown, whereas the full map of residues from 333 to 527 was shown in **Supplementary** Fig.7. **c-d**, Crystal structure of CC12.1 interacting with SARS-CoV-2 RBD modified from PDB: 6XC2 ^24^. CC12.1 heavy chain and light chain were colored in dark green and light green respectively. Key binding residues N501, K417, N460 and D420 on RBD were highlighted in orange. **e-f**, Representative neutralization curves of CC12.1, eCC12.1, eCC12.1.6, and eCC12.1.7 against SARS-CoV-2 **e** D420K and **f** N460H. Antibodies were colored according to the key. Error bars represent standard deviation. Data were representative of at least two independent experiments. **g**, Neutralization potency of parental CC12.1 and eCC12.1 variants against wild type SARS-CoV-2 virus and D420K, N460H, N460P, N460A mutant viruses.

**Fig. 7.**
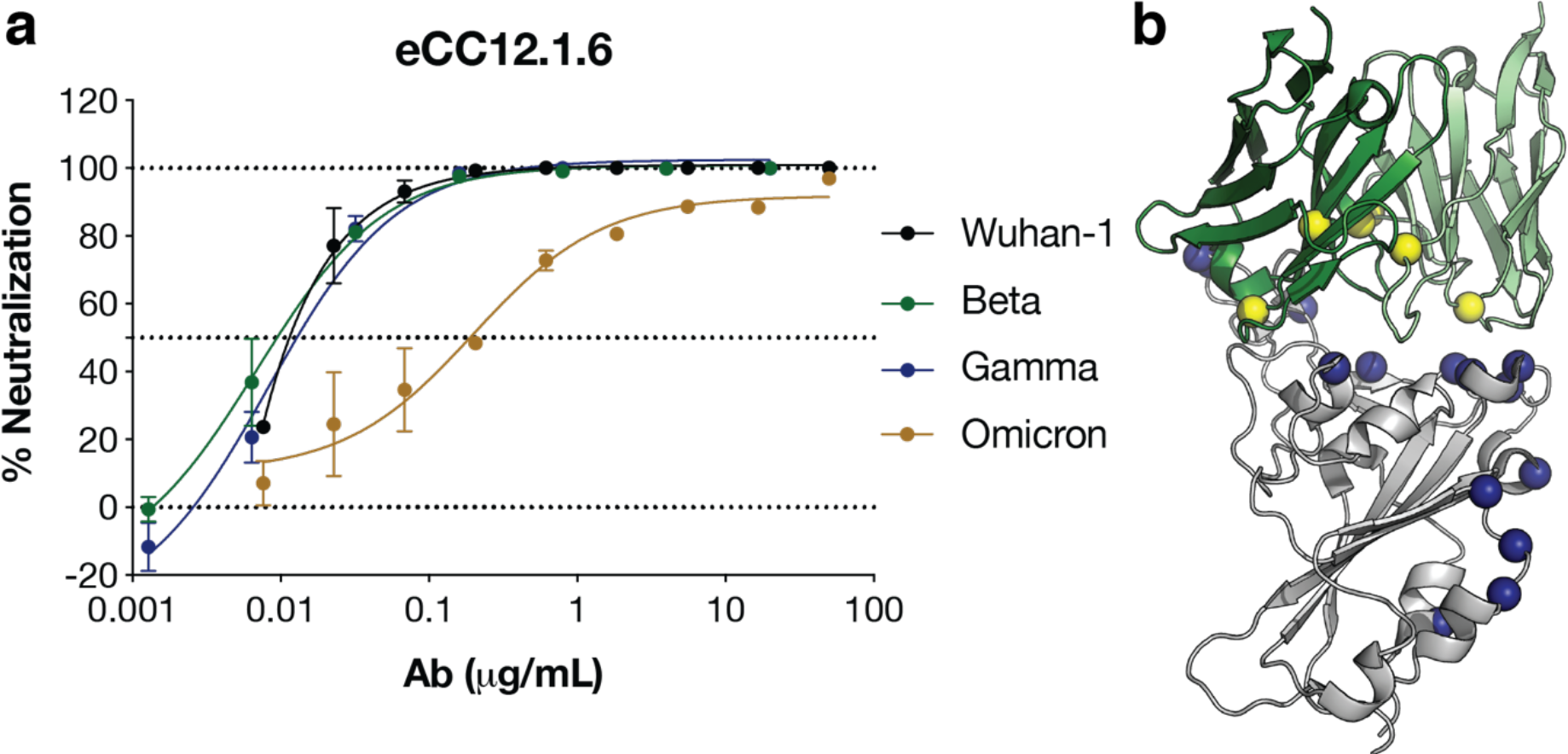
Neutralization and interaction of eCC12.1.6 against the Omicron variant. **a**, Representative neutralization curves of eCC12.1.6 against SARS-CoV-2 VOCs. Wildtype SARS- CoV-2 and VOCs were colored according to the key. Error bars represent standard deviation. **b**, Structure of eCC12.1.6 interacting with Omicron RBD modified from PDB: 6XC2^24^. eCC12.1.6 heavy chain and light chain were colored in dark green and light green respectively. Antibody mutant residues were highlighted in yellow spheres while Omicron RBD mutant residues were highlighted in navy blue spheres.

**Fig. 6.**
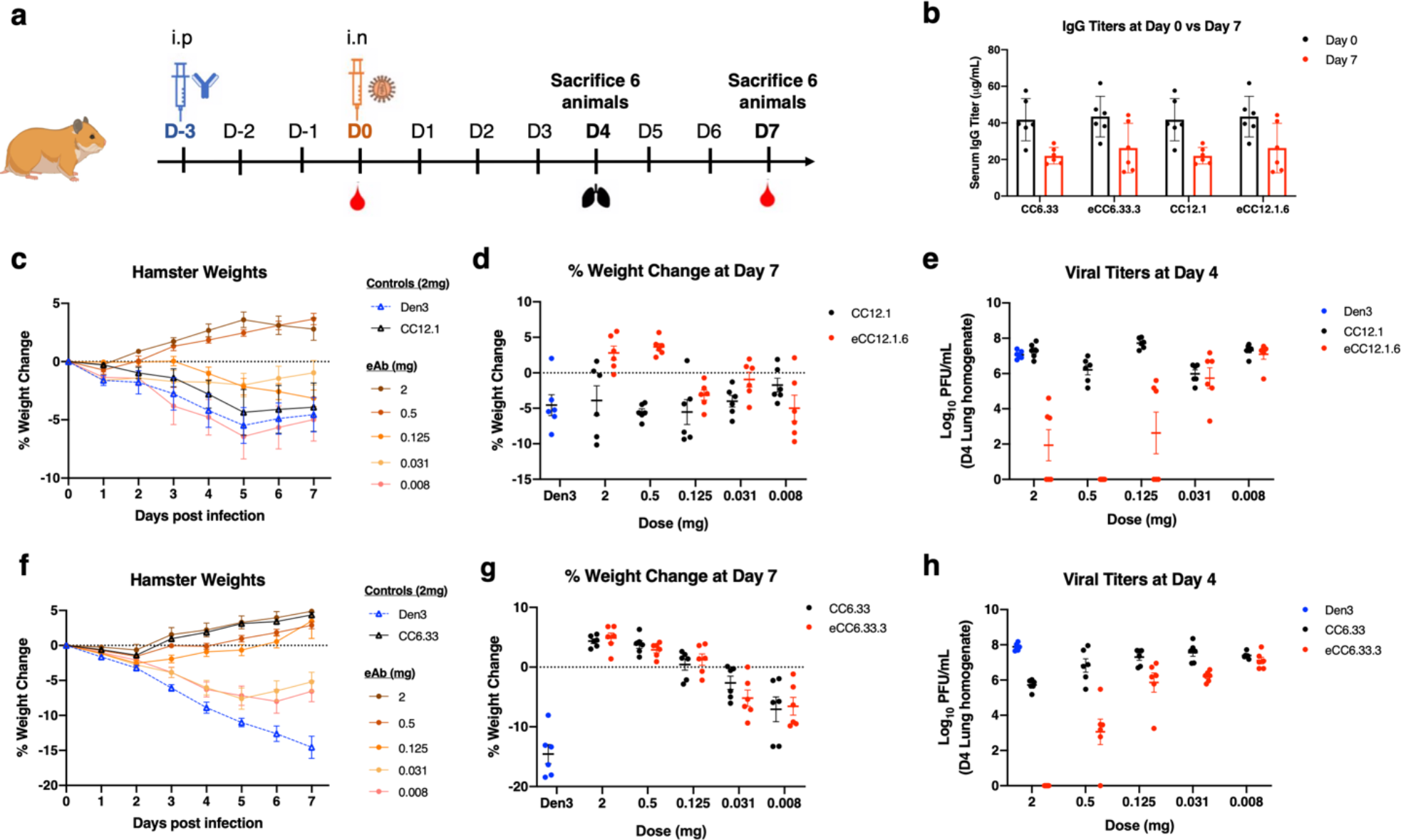
Protection of hamsters against SARS-CoV-2 challenge by parental and engineered nAbs. **a**, Overview of study design. Animal groups administered CC12.1 or eCC12.1.6 were challenged with 1 × 10^5^ PFU of SARS-CoV-2 (hCoV-19/South Africa/KRISP-EC-K005321/2020), those administered with CC6.33 or eCC6.33.3 were challenged with 1 × 10^5^ PFU of SARS-CoV-2 (USA-WA1/2020). **b**, Serum antibody concentration at time of infection (Day 0) versus the time of sacrifice (Day 7). **c**, Effect of CC12.1 vs eCC12.1.6 on weight loss in prophylaxis groups. **d**, Percent weight change for all groups from **c** on day 7 post infection. **e**, Viral load 4 days post infection as quantitated by live virus plaque assay on Vero E6 cells from lung tissue homogenate. Error bars represent geometric standard deviations of the geometric mean. **f**, Effect of CC6.33 vs eCC6.33.3 on weight loss in prophylaxis groups. **g**, Representative percent weight change for all groups from **f** on day 7 post infection. **h**, viral load, detected as shown in **e**. Statistical significance (p < 0.05) of groups in comparison to the Den3 IgG control group were calculated by Ordinary One-Way ANOVA test using Graph Pad Prism 8.0. Error bars represent group average with standard error of the mean.

### Neutralizing activity of engineered antibodies

The parental and engineered nAbs were tested for *in vitro* neutralization of SARS-CoV-1 and SARS-CoV-2 pseudotyped viruses^23^ to investigate the relationship between improved binding affinity and *in vitro* neutralization potency. The results were different for the 3 antibodies. All eCC6.33 variants showed improved neutralization potency, with the IC50 improving from 228 ng/mL to around 10 ng/mL for SARS-CoV-2 (Fig.3b) and from 2.27 µg/mL to around 20 ng/mL for SARS-CoV-1. All eCC6.33 variants achieved 100% inhibition while the parental CC6.33 had a maximum percent neutralization (MPN) of only around 80% (Supplementary Table 3). In stark contrast and despite comparable affinity improvements to eCC6.33 variants, eCC12.1 and eCC6.30 variants showed no significant improvement in neutralization potency relative to the parent Abs. To further investigate, we produced the parental and engineered nAbs as molecular Fabs to evaluate neutralization in a monovalent format. In all cases the engineered Fabs neutralized more potently than the parental Fabs, with parental CC12.1 and CC6.33 Fabs failing to neutralize at concentrations of 50 µg/mL (Fig.3c-e). As a control, we also tested our enAbs against authentic SARS-CoV-2 and observed similar neutralization activity to the pseudotyped virus (Fig.3f), consistent with our previous observations. Taken together, this data suggest that, in this system, increases in binding affinity translate to increases in the *in vitro* neutralization potency until a “threshold” IC50 around 10 ng/mL is reached, at which point further increases in binding affinity do not appear to affect the *in vitro* neutralization function of the antibody. Furthermore, this apparent affinity required to reach this neutralization threshold is lowered by the bivalent binding of an IgG. However, a Fab can neutralize the virus provided the monovalent affinity is sufficiently high, indicating inter- or intra- spike cross linking may help but is not necessary for these nAbs to neutralize (Fig.3c and Fig.3e).

We next sought to investigate how the evolving viral diversity of SARS-CoV-2 variants impacts the binding and the neutralization function of our parental and select engineered nAbs. We first measured the neutralization potency of our nAbs against VOCs with full-spike mutations including Alpha (B.1.1.7, originating in UK), Beta (B.1.351, originating in South Africa)^30^, Gamma (P.1, originating in Brazil)^31^, Kappa (B.1.617.1, originating in India) and Delta (B.1.617.2, originating in India)^32, 34^ variants as well as single mutations on RBD (Fig.4a).

CC12.1 neutralized Alpha, Delta and Kappa VOCs with an IC50 comparable to Wuhan-1 virus, however, Beta and Gamma VOCs completely escaped from this nAb (Fig.4b,c). Analysis of the individual variants found that K417N (from Beta VOC) and K417T (from Gamma VOC) facilitated this escape, consistent with the previous observation that most VH3-53-class nAbs are sensitive to these mutations^14, 35, 36^. K417 falls in the middle of the CC12.1 crystallographically defined epitope and makes hydrogen bonding or packing interactions with heavy chain residues H33, Y52, D95 and D97, as well as light chain residues N92 and K97^24^. Importantly, eCC12.1.4 and eCC12.1.7 neutralized all the VOCs, including Beta and Delta containing the K417N/T mutations (Fig.4c). We measured neutralization of all 12 eCC12.1 antibodies against Beta and Gamma as well as the single mutation variants, where we found 11 out of 12 mAbs neutralized the Gamma lineage and 9 out of 12 neutralized the Beta lineage (Supplementary Fig.6a). Mutational analysis of the 9 enAbs that reacted against both VOCs found a broad assortment of mutations that had been selected across the different antibodies (Supplementary Fig.6a), suggesting that there are multiple ways to compensate for the loss of the K417 interaction.

CC6.30 and eCC6.30.2 were effective against the Alpha variant, showed significantly reduced function against the Delta variant and were completely unable to neutralize the Beta, Gamma or Kappa variants (Fig.4b,c). When tested against pseudoviruses containing the individual mutations found in these variants, CC6.30 showed complete loss of neutralization against the single L452R, E484K and E484Q variants. eCC6.30.2 retained modest functionality against L452R and E484Q, while the E484K variant still facilitated complete viral escape. This data was consistent with our structural analysis, with both E484 and L452 making extensive interactions with CC6.30 (Fig.1c,d).

The parental CC6.33 and eCC6.33.8 were effective at neutralizing all VOCs tested with similar IC50s to the original Wuhan-1 SARS-CoV-2, consistent with the observation that CC6.33 recognizes the conserved class 3 epitope^5, 28^ distal from the mutations in these viruses (Fig.2b, Fig.4b,c, and Supplementary Fig.6b). REGN10987, another class 3 antibody^5, 27^ retained similar potency for all VOCs tested (Supplementary Fig.6b).

To systematically investigate the relationship between binding affinity and neutralization for VOCs across our collection of parental and engineered nAbs, a panel of monomeric RBD variants were expressed for SPR analysis. Overall, the RBD binding affinities and the off-rate correlated well with the *in vitro* neutralization data (Fig.4d-f). Mutations that completely abrogated neutralization usually showed a complete loss of binding by SPR or weak reactivity that could not be fit to a simple kinetics model. Of particular note were the Beta and Gamma RBDs binding to the CC12.1 variants. Parental CC12.1 bound Wuhan-1 RBD with an affinity of 6 nM, and had a complete loss of both binding and neutralization to both Beta and Gamma RBDs. In contrast, affinity matured eCC12.1.4 bound Wuhan-1 with an affinity of 286 pM, and although the mutations in Beta and Gamma reduced binding affinity by 182- fold and 43- fold (Supplementary Table 4), respectively, eCC12.1.4 was still able to neutralize both VOCs (Fig.4c). These data indicate that enhanced nAb affinity for the target antigen helps to offset the affinity losses resulting from viral mutations within the nAb paratope, allowing the nAb to maintain sufficient affinity for neutralization.

### Mapping RBD escape mutations for CC12.1 and eCC12.1.4

We next asked whether engineering high affinity nAbs reduces the pathways for viral escape compared with the parental nAbs or if they simply shifted the escape mutations to other positions. Deep mutational scanning libraries of RBD were generated and used to determine the mutations on RBD that prevented CC12.1 and eCC12.1.4 from blocking ACE2 binding *in vitro* (Supplementary Fig.7a)^36^. 94.5% (2250/2380) of all possible single mutations were scanned (Supplementary Table 5). Consistent with previous reports^36^ and our neutralization screening, CC12.1 is vulnerable to multiple mutations at K417 with a false discovery rate (FDR) below 0.1 for K417N/T but eCC12.1.4 is able to accommodate all mutations at K417 (Fig.5a,b, and Supplementary Fig.7b). We also detected multiple mutations at position D420 and N460 that confer escape from the parental CC12.1 (Fig.5a,b). Alanine scanning and pseudovirus escape mutations had identified these D420 and N460 residues as important for public VH3-53 SARS-CoV-2 antibodies^35, 37, 38^, but structural analysis shows these two positions on the periphery of the CC12.1 epitope and making relatively insignificant contacts to the antibody (Fig.5c,d).

Pseudoviruses containing the individual D420K, N460H, N460P, and N460A mutations, identified as potential escape mutations in the deep mutational scanning, were produced and evaluated to determine if parental CC12.1 and several eCC12.1 variants were sensitive to these mutations in a neutralization assay. In agreement with RBD library screening, a D420K substitution completely disrupted CC12.1 neutralization, while substitutions at the N460 residue significantly decreased its neutralization potency by 12- to 246-fold (Fig.5e-g). By contrast, although D420K and N460H were identified as potential escape mutations against eCC12.1.4, neutralization potency was reduced by a more modest 8-fold against D420K and remained insensitive to a N460H substitution (Fig.5d,f). These data suggest that increasing the affinity of SARS-CoV-2 nAbs restricts the potential escape mutations that can arise in RBD, rather than just altering the critical nAb contacts and shifting the escape mutations to a comparable number of different positions and/or mutations. This is particularly important in the context of developing antiviral antibodies where viral escape is a serious and constant threat.

### In vivo protection

*In vitro* analysis of the enAbs suggested that the increased affinity provided functional improvements for two of the three candidates. eCC12.1 variants were better able to overcome mutations that are emerging in the VOCs and eCC6.33 variants had increased neutralization potency and higher MPN compared to CC6.33. The improvements to CC6.30 were less clear cut and the parental antibody was found to have poor pharmacokinetics in hamsters and so was not pursued further *in vivo* (data not shown). A series of experiments were designed to compare parental and engineered nAbs in the Golden hamster model of COVID-19 infection. The experimental design for passive transfer studies is shown in Fig.6a. Groups of six hamsters were prophylactically treated with serially diluted doses of antibody starting at 2 mg per animal to 8 µg per animal via intraperitoneal (i.p.) injection 72 hours before intranasal challenge with SARS-CoV- 2 at a dose of 1 x 10^5^ plaque-forming units (PFU). A group receiving 2 mg doses of an irrelevant human mAb against dengue virus (Den3) was used as a control for each experiment. All hamsters were monitored daily for weight loss as a measure of disease^39^ and serum was collected from each animal to determine antibody titer at the time of viral challenge (D0) compared to the time of sacrifice (D7) (Fig.6b). Hamsters have been shown to clear SARS-CoV-2 infection after 7 days, so a replicate of the original experiment was performed in which the groups of hamsters were euthanized four days post infection (D4) and lung tissue was collected to quantify lung viral titers. In addition to collecting serum at D0 and D7 to measure nAb titers, the half-life of the parental and several engineered versions were assessed to try to find engineered nAbs that closely matched the bioavailability of the parental versions to allow a comparison of the two. Ultimately, eCC6.33.3 and eCC12.1.6 were selected to compare to the parental nAbs.

We first tested the ability of CC12.1 and eCC12.1.6 to protect against challenge from the Beta VOC, as the *in vitro* data showed that eCC12.1.6 effectively neutralized the variant (Supplementary Fig.6b). The prophylactic protection experiment described above was done using Beta (20H/501Y.V2) SARS-CoV-2. Consistent with our *in vitro* neutralization data, eCC12.1.6 exhibited a dose-dependent protective response both in terms of weight loss and lung viral titers (Fig.6c-e). Parental CC12.1 showed no protection compared to the Den3 control group. We also assessed eCC12.1.6 against the original SARS-CoV-2 (USA/WA1/2020) using the same groups (Supplementary Fig.8) and the weight loss trend was nearly identical to that of the Beta variant, albeit the Beta variant showed a lower overall percentage of weight loss (Supplementary Fig.8e).

The second protection experiment was designed to test whether the increased *in vitro* neutralization potency and maximum neutralization percentage of eCC6.33.3 relative to the parental nAb provided enhanced protection. CC6.33 (*in vitro* IC50 = 0.228 µg/mL and MPN of 81%) was compared with eCC6.33.3 (*in vitro* IC50 = 0.008 µg/mL and MPN of 100%) for protective efficacy against challenge from the original SARS-CoV-2 (USA/WA1/2020) (Supplementary Fig.9). Following viral challenge, animals that received either the parental or engineered nAb, including the groups that received the 8 µg dose, showed a statistically significant reduction in weight loss compared to the group receiving the Den3 (Fig.6f,g). The protection from weight loss correlated with the amount of nAb the animals received, and there was no apparent difference in efficacy between the parental and engineered nAbs. However, in contrast to the weight loss results, the enAbs showed superior ability to control viremia in the lung (Fig.6h). Animals that received the 2 mg dose of eCC6.33.3 had sterilizing immunity and the animals that received 500 µg dose of eCC6.33.3 had lung viral titers 5 logs lower than the Den3 control group. Animals that received the parental CC6.33 had viral titers only 2 logs lower than the Den3 group, protection that was comparable to the group receiving 125 µg of eCC6.33.3. It is unclear why there is a disconnect between lung viral titers and weight loss, however, eCC6.33.3 was clearly superior at controlling lung viral load. Broadly, the two experiments confirm that the affinity engineering of these nAbs provide a superior *in vivo* benefit predicted from *in vitro* analysis.

### Neutralization of the Omicron variant

During the preparation of this manuscript the Omicron VOC was reported. The VOCs characterized above contained 7 to 12 mutations in S compared to the original SARS-CoV-2, and at most contained 3 mutations in RBD (Beta and Gamma). In contrast, Omicron contains 30 mutations, 3 deletions and an insertion in S, with 15 of these mutations located in the RBD. The mutations in RBD are heavily concentrated across the class I, class II and class III neutralizing antibody epitopes and have been shown to reduce the neutralization efficacy of plasma from vaccinated and/or infected donors^40–43^. They also confer resistance or complete escape from the majority of clinical antibody candidates^43^.

Omicron completely escaped from the parental CC6.30 and all the eCC6.30 variants (Supplementary Fig.10). This was largely consistent with data from other VOCs demonstrating the importance of the E484 interaction for this antibody family. Omicron not only contains an E484A mutation, but also contains a Q493K mutation that removes a hydrogen bond between Q493 and R97H on CC6.30 variants. Similarly, the parental CC6.33 and all eCC6.33 variants were also unable to neutralize Omicron (Supplementary Fig.10). This observation was more unexpected. The only Omicron mutation immediately within the CC6.33 epitope is G339D that may introduce a clash with the CC6.33 CDRH3 backbone. Omicron also contains mutations S371L, S373P and S375F that could alter the conformation of an adjacent loop and prevent the N343 glycan from adopting the conformation observed in the CC6.33 bound structure. Finally, we observed that Omicron was resistant to parental CC12.1 and 11 of 12 eCC12.1 variants, however, eCC12.1.6 retained neutralizing activity against Omicron (IC50 of 0.20 µg/mL), albeit with approximately 25-fold reduced potency (Fig.7a, and Supplementary Fig.10). eCC12.1.6 was also the most effective antibody against Beta and Gamma VOCs, with comparable potency to the Wuhan-1 strain (Fig.7a). Analysis of the selected mutations in eCC12.1.6 compared to other eCC12.1 variants did find a unique S31W mutation in LCDR2 that is located in close proximity to the N501Y mutation in all VOCs as well as G466S, G496S, Q498R and Y505H that are present in Omicron (Fig.7b). It is also possible that the other mutations in the antibody that do not interact directly with RBD stabilize the CDR loops in a conformation that happened to be slightly more compatible with Omicron compared to the other eCC12.1 variants.

## DISCUSSION

Emerging SARS-CoV-2 VOCs have challenged vaccine-induced immunity and therapeutic mAbs, and functional nAbs with prophylactic and therapeutic efficacy against VOCs are desired. To this end, investigators have searched for nAbs that are largely resistant to viral antigenic variability by screening for nAbs that broadly neutralize SARS-CoV-2 VOCs and/or other sarbecoviruses. This effort has largely paralleled the work to identify broadly neutralizing HIV or influenza antibodies that target conserved epitopes on these antigenically variable viruses^44, 45^. Here, we explored an alternative and complementary strategy of engineering nAbs to have higher affinity for their target antigen through directed evolution and then investigated the relationship between binding affinity, *in vitro* neutralization potency and *in vivo* efficacy. Broadly, we found that monovalent binding affinity and *in vitro* neutralization potency are correlated, until the *in vitro* neutralization IC50 reaches a “threshold” (around 10 ng/mL for IgG) after which further affinity improvements did not translate to improvements in neutralization potency, at least for CC12.1 and CC6.30 nAbs that directly compete with ACE2 binding to the RBD. These affinity improvements do help to expand the breadth of antibody reactivity, allowing them to better neutralize VOCs that contain mutations in and around the antibody epitope. This was particularly evident with the eCC12.1 variants that are part of the shared VH3-53 lineage nAbs and are broadly susceptible to the K417T/N mutations found in Beta and Gamma VOCs^14, 36^. These mutations abrogated binding of the parental CC12.1 nAb, much like other reported VH3-53 nAbs; however, the higher affinity of the eCC12.1 variants for wild-type S-protein was able to compensate for lost the K417 contact allowing the nAb to maintain sufficient affinity that it could still potently neutralize the VOCs. Of note, eCC12.1 variants were affinity matured against the original SARS-CoV-2 RBD sequence and had no specific selective pressure to accommodate mutations in the VOCs. Importantly, our saturated mutagenesis screening showed that the affinity maturation restricted the number of potential escape mutations rather than just altering them to different positions. While increasing the affinity could restrict escape mutations, it did not abrogate them entirely, as eCC12.1 variants showed modest sensitivity to the D420K mutation, and all eCC6.30 variants were still unable to bind or neutralize Beta and Gamma VOCs with the E484K mutation. In the extreme case of the Omicron variant that contains so many mutations on the RBD, especially within the footprint of class 1 antibodies, there was still one CC12.1 variant that had significant neutralizing activity. This finding illustrates the value of affinity maturation in the context of natural infection in that the generation of a diverse set of related antibodies, as was done here in vitro, will likely generate some antibodies able to bind to and act against many different viral variants, including those with multiple mutations as for Omicron (and see below). It is possible that part of the reason so many clinical antibody candidates failed against Omicron is that most were selected shortly after COVID infection before much affinity maturation had occurred^46^.

The engineering of higher affinity enAbs also improved protective efficacy *in vivo*. The protection data also show that eCC12.1.6 was effective at preventing infection from the Beta variant while CC12.1 was not, consistent with the *in vitro* neutralization results. eCC6.33.3 provided significantly enhanced viral control with sterilizing immunity at the 2 mg dose and ∼100,000 fold reduction in lung viral titers at the 500 µg dose compared to the Den3 control. A plausible explanation for this enhanced control is the improvement in neutralization plateau—CC6.33 only achieves ∼80% MPN whereas eCC6.33 variants plateau at 100% neutralization. While the results were expected, it was important to formally demonstrate that the affinity/neutralization improvements translated to enhanced protection *in vivo*. We also note that, while not the objective of the experiment, the engineering produces a broad spectrum of related nAbs with different biochemical properties and this resulted in an eCC6.30 variant with significantly better pharmacokinetics compared to the parental CC6.30. Taken together, this data strongly suggests the benefit of engineering anti-viral nAbs for improved affinity to their target antigen, with an important benefit being the restriction of potential escape pathways for the virus.

The data also emphasizes the importance of affinity maturation in vaccine-induced responses to maximize the opportunities for protection against viral variants. Longitudinal studies have shown that mAbs isolated from later time points have more somatic mutation compared to clonally related variants from earlier time points, and this additional somatic mutation can increase affinity, potency and expand breadth^38, 47^. The enAb selections demonstrated that there are numerous solutions from a single starting point that can produce antibodies with increased binding affinity (Supplementary Table 3). This situation is similar to a vaccine-expanded B cell lineage, that undergoes slightly different maturation trajectories. Despite achieving similar final results against the target antigen, the different members within the expanded lineage can have different levels of cross-reactivity to viral variants. We also note that the binding kinetics of our enAbs are consistent with what can be achieved through multiple vaccinations with a soluble protein antigen^48^. These data support the current approach of boosting with the original SARS-CoV-2 vaccine formulation, as continued expansion of B cell lineages and further improvements in affinity can help expand the response to cover new VOCs. It also will help to expand the pool of memory B cells that may be cross-reactive to new vaccine formulations when they are eventually developed.

In summary, we demonstrate that increasing antibody affinity can result in improved neutralization breadth and potency, and these improvements can enhance the therapeutic antibody function.

## Materials and Methods

### Syrian hamsters

Golden Syrian hamsters were provided by Charles River Laboratories (CRL:LVG(SYR)) and housed at the Scripps Research Institute. Male 12–13-week-old hamsters were infused with antibodies intraperitoneally as described previously^23^. The Scripps Research Institutional Animal Care and Use Committee (IACUC) approved all experimental procedures involving all the animals in accordance with Protocol #20-0003.

### Cell lines

Saccharomyces cerevisiae YVH10 cells (ATCC) were used in antibody library generation and FACS sort. Hela-hACE2 cells^23^ were used in pseudovirus neutralization assay. Saccharomyces cerevisiae EBY100 cells (ATCC) were used in RBD mutant library generation and FACS sort. Human HEK293T cells (ATCC) were used for pseudovirus production. FreeStyle HEK293 cells (ThermoFisher) were used for recombinant S protein production. Expi293F cells (ThermoFisher) were used for monoclonal antibody and recombinant RBD production. Vero-E6 cells (ATCC) were used for live virus plaque assay.

### Recombinant S and RBD production

SARS-CoV-1 (Genbank AAP13567) or SARS-CoV-2 (Genbank MN908947) S proteins were transiently expressed in Freestyle 293F system (ThermoFisher) whereas RBD proteins were expressed in the Expi293 system (ThermoFisher). In brief, S expression plasmids were cotransfected with 40K PEI (1 mg/mL) at a ratio of 1:3. After incubation for 30 min at RT, transfection mixture was added to Freestyle 293F cells at a density of approximately 1 million cells/mL. RBD plasmids with His-Avitag were cotransfected with FectoPRO (Polyplus 116-010). SARS-CoV-2 RBD mutant plasmids were generated by Quikchange site-directed mutagenesis according to manufacturer’s instructions (Agilent, 210513). Biotinylated proteins were made by co-transfecting Avitagged RBD plasmids with a BirA expression plasmid and into Expi293 cells using FectoPRO. After incubation for 10 min at RT, transfection mixture was added to Expi293 cells at a density of ∼ 3 million cells/mL. After 24h of transfection, cells were fed with D-(+)- glucose solution and 300 mM of sterile sodium valproic acid solution. After 5 days of transfection, supernatants were harvested and filtered with a 0.22 µm membrane.The His-tagged proteins were purified with the HisPur Ni-NTA Resin (Thermo Fisher, 88222). After three columns of washing with 25 mM Imidazole (pH 7.4), proteins were eluted with an elution buffer (250 mM Imidazole, pH 7.4) at slow gravity speed (∼4 sec/drop). Eluted proteins were buffer exchanged and concentrated with PBS using Amicon tubes (Millipore). The proteins were further purified by size exclusion chromatography (SEC) using Superdex 200 (GE Healthcare). The selected fractions were pooled and concentrated.

For cryoEM, a stabilized version of CoV-2 S protein, HPM7 (hexaproline mutant 7) was used. The design, expression and purification has been described previously^14^. Briefly, the S protein is stabilized with 6 engineered proline residues^29^ and an interprotomer disulfide between residues 705 and 883 of S2. HPM7 was expressed in HEK293F, and purified using a C-terminal 2x StrepTag followed by size exclusion chromatography to isolate trimers.

### Antibody production and purification

Monoclonal antibody was transiently expressed in the Expi293 system (ThermoFisher, A14635). In brief, antibody HC and LC plasmids were co-transfected at a ratio of 1:2.5 with transfection reagent FectoPRO (Polyplus 116-010). After 24 h of transfection, 300 mM of sterile sodium valproic acid solution (Sigma-Aldrich, P4543) and 45% D-(+)- glucose solution (Sigma Aldrich, G8769-100ML) were added to feed cells. After 4-5 days of transfection, supernatants were collected, sterile-filtered (0.22 µm), and IgG was purified with Protein A sepharose beads (GE Healthcare 17-5280-04).

### Expression and purification of Fab

The CC6.33, eCC6.33.3, CC6.30, eCC6.30.2 Fabs were purified by digesting IgG using Fab digestion kit (ThermoFisher, 44985) according to manufacturer’s instructions. After digestion, Fc fragments and undigested IgG were removed from binding to the protein A beads (GE Healthcare). CC12.1 and eCC12.1.7 Fab fragments were generated by introducing a stop codon after the CH1 region in heavy chain expression plasmids. After transfection, Fabs were purified using Capto L beads (Cytiva, 17547801).

### CryoEM data collection and processing

To form the antibody-spike complexes, CC6.30 Fab or CC6.33 IgG was incubated with HPM7 spike trimer for approximately 15 minutes. A 9:2 molar ratio of CC6.30 Fab to spike or 1:2 molar ratio of CC6.33 IgG to spike was used. Samples were concentrated to about 1.7 mg/mL, n-dodecyl-β-D-maltopyranoside (DDM) was added to a final concentration of 0.06 mM, and the mixture and applied to plasma cleaned Quantifoil 1.2/1.3 400 copper mesh, holey carbon grids. Grids were vitrified using the Vitrobot Mark IV system (Thermo Fisher) set to 4°C, 100% humidity, with a blot time of 3 seconds.

Data were collected on a Thermo Fisher Titan Krios equipped with a Gatan K2 Summit direct electron detector. Detailed data collection statistics are summarized in Supplementary Table 1 and 2. Movie micrographs were motion corrected and dose weighted using MotionCorr2^49^. Aligned micrographs were imported into cryoSPARC version 3.2^50^ and CTF corrections were performed using the Patch CTF application in cryoSPARC. Automated particle picking, particle extraction, and initial 2D classifications were performed in cryoSPARC. Particles belonging to selected 2D classes were then imported into Relion 3.1^51^. Interactive rounds of 3D classification and refinement, and CTF refinements were performed for each dataset. To improve resolution of the antibody epitope and paratope, the best refinement from each dataset (Spike with 2 Fabs bound for both CC6.30 and CC6.33) was subjected to C3 symmetry expansion and focused classifications, using a spherical mask around the expected Fab/RBD region of a single protomer and Relion 3D classification without alignments. Particles containing density for Fab and RBD in the region of interest were imported into cryoSPARC. Signal outside of the RBD and Fab Fv was subtracted, and the subtracted particles were subjected to cryoSPARC local refinement (Supplementary Fig.1-2). The non-symmetry expanded particles from the best Relion global refinements were imported into cryoSPARC and subjected to a final round of non-uniform refinement ^52^. Additionally, the CC6.33 dataset contained thousands of ligand-free HMP7 Spike particles which were also imported into cryoSPARC for final non-uniform refinement, with or without symmetry (Figure S2C and S2D). Final Fourier shell correlation resolution estimates for all maps, along with EMDB deposition codes, can be found in Supplementary Table 1 and 2.

### CryoEM model building

Homology models of the Fab variable regions of CC6.30 and CC6.33 were generated using ABodyBuilder^53^ and fitted into the respective local refinement maps using UCSF Chimera^54^. Coordinates for RBD with complete ridge were taken from PDB 7byr. The RBD:Fv models were subjected to interactive cycles of manual and automated refinement using Coot 0.9^55^ and Rosetta^56^. Once a high map-to-model agreement was reaching, as measured by EMRinger^57^, and geometries were optimized, as judged by MolProbity^58^, the models were fit into the non-uniform refinement full trimer maps and combined with a Spike model refined into the ligand-free map (PDB 6vxx was used as the initial model and HPM7 mutations were added manually in Coot following iterative rounds of Rosetta relaxed refinement and Coot manual editing). The resulting Fv:trimer models were refined in Rosetta. The Phenix software suite^59^ was used for structure validation, and for editing and preparation of PDB files for deposition. Final refinement statistics and PDB deposition codes for generated models can be found in Supplementary Table 1 and 2.

### Antibody library generation

CC12.1, CC6.30 and CC6.33 heavy chain and light chain affinity maturation libraries were synthesized as Oligopools (Integrated DNA Technologies). Mutations were included in the CDR loops and the CDR1/2/3 mini-libraries were assembled into combinatorial heavy chain and light chain libraries as previously described^33^. The libraries were displayed on the surface of yeast as molecular Fab using the yeast display vector pYDSI containing the bidirectional Gal1-10 promoter. The heavy chain contains a C-terminal V5 epitope tag and the light chain contains a C-terminal C-myc epitope tag to assess the amount of Fab displayed on the surface of the yeast. The HC library was generated by cloning the HC CDR1/2/3 library into the vector containing the wildtype light chain by homologous recombination, and the LC library was generated by doing the inverse. The combinatorial H/L library was generated by amplifying the HC and LC sequences with primers overlapping in the Gal1-10 promoter. The recovered Gal-HC and Gal-LC fragments were ligated via Gibson assembly and amplified. The resulting LC-Gal1-10-HC product was cloned into an empty pYDSI vector by homologous recombination^33^.

### Yeast library labeling and sorting

After yeast transformation, yeast cells were passaged 1:20 the following day. Cells were then induced at OD600 = 1.0 overnight at 30°C in SGCAA induction medium (20 g galactose, 1 g glucose, 6.7 g yeast nitrogen base without amino acid, 5 g bacto casamino acids, 5.4 g Na2HPO4, 8.56 g NaH2PO4!”2O, 8.56 mg uracil in 1 L deionized water, pH 6.5, and sterilize by filtration). For each library, in the first round of selection, 5 x 10^7^ of yeast cells were stained per sample. In the second to final round of selection, 1 x 10^7^ cells were stained. Yeast cells were firstly spun down and washed with PBSA (PBS + 1% BSA), then incubated with biotinylated SARS-CoV-2 RBD or S or HEK cell membrane protein at several non-depleting concentrations respectively for at least 30 min at 4°C. After washing, yeast cells were stained with FITC-conjugated chicken anti-C-Myc antibody (Immunology Consultants Laboratory, CMYC-45F), AF405-conjugated anti-V5 antibody (made in house), and streptavidin-APC (Invitrogen, SA1005) in 1:100 dilution for 20 min at 4 °C. After washing, yeast cells were resuspended in 1 mL of PBSA and loaded on BD FACSMelody cell sorter. Top 5-10% of cells with high binding activity to a certain SARS-CoV-2 RBD labeling concentration were sorted and spun down. Sorted cells were expanded in 2 mL of synthetic drop-out medium without tryptophan (Sunrise, 1709-500) supplemented with 1% Penicillin/Streptomycin (Corning, 30-002-C) at 30°C overnight.

### Size exclusion chromatography analysis

The antibodies were analyzed by size exclusion chromatography using the 1260 Infinity II (Agilent). 15 uL of each antibody at 2 mg/mL was injected into the TSKgel SuperSW mAb column (Tosoh) with the flow rate of 1 mL/min.

### Pseudovirus neutralization assay

Pseudovirus was generated as described previously^23^. In brief, MLV gag/pol backbone (Addgene, 14887), MLV-CMV-Luciferase plasmid (Addgene, 170575), and SARS-CoV-2-d18 (Genbank MN908947) or SARS-CoV-1-d28 (Genbank AAP13567) or SARS-CoV-2 VOC spike plasmid were incubated with transfection reagent Lipofectamine 2000 (Thermo Fisher, 11668027) following manufacturer’s instructions for 20 min at RT. Full-spike mutations were introduced by overlapping extension polymerase chain reaction (PCR) to generate mutated spikes of circulating SARS-CoV- 2 VOC, i.e., B.1.1.7, B.1.351, P.1, B.1.617.1, B.1.617.2, and B.1.1.529. Then the mixture was transferred onto HEK 293T cells (ATCC, CRL-3216) in a 10 cm^2^ culture dish (Corning, 430293). After 12-16 h of transfection, culture medium was gently removed, fresh DMEM medium was added onto the culture dish. Supernatants containing pseudovirus were harvested after 48 h post transfection and frozen at -80 °C for long term storage.

In the neutralization assay, antibody samples were serially diluted with complete DMEM medium (Corning, 15-013-CV) containing 10% FBS (Omega Scientific, FB-02), 2 mM L-Glutamine (Corning, 25-005-Cl), and 100 U/mL of Penicillin/Streptomycin (Corning, 30-002-C). 25 µL/well of diluted samples were then incubated with 25 µL/well of pseudotyped virus for 1 h at 37 °C in 96-well half-area plates (Corning, 3688). After that, 50 µL of Hela-hACE2 cells at 10,000 cells/well with 20 µg/mL of Dextran were added onto each well of the plates. After 48 h of incubation, cell culture medium was removed, luciferase lysis buffer (25 mM Gly-Gly pH 7.8, 15 mM MgSO4, 4 mM EGTA, 1% Triton X-100) was added onto cells. Luciferase activity was measured by BrightGlo substrate (Promega, PR-E2620) according to the manufacturer’s instructions. mAbs were tested in duplicate wells and independently repeated at least twice. Neutralization IC50 values were calculated using “One-Site Fit LogIC50” regression in GraphPad Prism 8.0.

### Authentic SARS-CoV-2 neutralization assay

Vero-E6 cells were seeded in 96-well half-well plates at approximately 8000 cells/well in a total volume of 50 µL complete DMEM medium the day prior to the addition of antibody and virus mixture. The virus (500 plaque forming units/well) and antibodies were mixed, incubated for 30 minutes and added to the cells. The transduced cells were incubated at 37°C for 24 hours. Each treatment was tested in duplicate. The medium was removed and disposed of appropriately. Cells were fixed by immersing the plate into 4% formaldehyde for 1 hour before washing 3 times with phosphate buffered saline (PBS). The plate was then either stored at 4°C or gently shaken for 30 minutes with 100 µL/well of permeabilization buffer (PBS with 1% Triton-X). All solutions were removed, then 100 µl of 3% bovine serum albumin (BSA) was added, followed by room temperature (RT) incubation at 2 hours.

A mix of primary antibodies consisting of CC6.29, CC6.33, CC6.36, CC12.23, CC12.25^23^ in equal amounts for detection. The primary antibody mixture was diluted in PBS/1% BSA to a final concentration of 2 µg/ml. The blocking solution was removed and washed thoroughly with wash buffer (PBS with 0.1% Tween-20). The primary antibody mixture, 50 µl/well, was incubated with the cells for 2 hours at RT. The plates were washed 3 times with wash buffer.

Peroxidase AffiniPure Goat Anti-Human IgG (H+L) secondary antibody (Jackson ImmunoResearch, 109-035-088) diluted to 0.5 µg/mLl in PBS/1% BSA was added at 50 µL/well and incubated for 2 hours at RT. The plates were washed 6 times with wash buffer. HRP substrate (Roche, 11582950001) was freshly prepared as follows: Solution A was added to Solution B in a 100:1 ratio and stirred for 15 minutes at RT. The substrate was added at 50 µL/well and chemiluminescence was measured in a microplate luminescence reader (BioTek, Synergy 2).

The following method was used to calculate the percentage neutralization of SARS-CoV-2. First, we plotted a standard curve of serially diluted virus (3000, 1000, 333, 111, 37, 12, 4, 1 PFU) versus RLU using four-parameter logistic regression (GraphPad Prism 8.0).

### Recombinant protein ELISA

6x-His tag antibodies (Invitrogen, MA1-21315) were coated at 2 µg/mL in PBS onto 96-well half-area high binding plates (Corning, 3690) overnight at 4°C. After washing and blocking, 1 µg/mL of his tagged recombinant SARS-CoV-2 (or SARS-CoV-1) RBD or S proteins were diluted in PBS with 1% BSA and incubated for 1 h at RT. After washing, serially diluted antibodies were added in plates and incubated for 1 h at RT. After washing, alkaline phosphatase-conjugated goat anti-human IgG Fcγ secondary antibody (Jackson ImmunoResearch, 109-055-008) was added in 1:1000 dilution and incubated for 1 h at RT. After final wash, phosphatase substrate (Sigma-Aldrich, S0942- 200TAB) was added into each well. Absorption was measured at 405 nm.

### Polyspecificity reagent (PSR) ELISA

Solubilized CHO cell membrane proteins (SMP) were made in house. SMP, human insulin (Sigma-Aldrich, I2643), single strand DNA (Sigma-Aldrich, D8899) were coated onto 96-well half-area high-binding ELISA plates (Corning, 3690) at 5 µg/mL in PBS overnight at 4°C. After washing, plates were blocked with PBS/3% BSA for 1 h at RT. Antibody samples were diluted at 100 µg/mL in 1% BSA with serial dilution and then added in plates and incubated for 1 h at RT. After washing, alkaline phosphatase-conjugated goat anti-human IgG Fcγ secondary antibody (Jackson ImmunoResearch, 109-055-008) was added in 1:1000 dilution and incubated for 1h at RT. After final wash, phosphatase substrate (Sigma-Aldrich, S0942-200TAB) was added into each well. Absorption was measured at 405 nm.

### HEp2 epithelial cell polyreactive assay

Reactivity to human epithelial type 2 (HEp2) cells was determined by indirect immunofluorescence on HEp2 slides (Hemagen, 902360) according to manufacturer’s instructions. Briefly, mAbs were diluted at 100 µg/mL in PBS respectively and then incubated onto immobilized HEp2 slides for 30 min at RT. After washing, one drop of FITC-conjugated goat anti-human IgG antibody was added onto each well and incubated in the dark for 15-30 min at RT. After washing with PBS, a cover slide was added to HEp2 cells with glycerol and the slide was photographed on a Nikon fluorescence microscope to detect GFP. All panels were shown at magnification 40x.

### Surface plasmon resonance methods

SPR measurements were carried out on a Biacore 8K instrument at 25°C. All experiments were carried out with a flow rate of 30 µL/min in a mobile phase of HBS-EP [0.01 M HEPES (pH 7.4), 0.15 M NaCl, 3 mM EDTA, 0.0005% (v/v) Surfactant P20]. Anti-Human IgG (Fc) antibody (Cytiva) was immobilized to a density ∼7000-10000 RU via standard NHS/EDC coupling to a Series S CM- 5 (Cytiva) sensor chip. A reference surface was generated through the same method.

For conventional kinetic/dose-response, listed antibodies were captured to 50-100 RU via Fc-capture on the active flow cell prior to analyte injection. A concentration series of SARS-CoV-2 RBD was injected across the antibody and control surface for 2 min, followed by a 5 min dissociation phase using a multi-cycle method. Regeneration of the surface in between injections of SARS-CoV- 2 RBD was achieved by a single, 120s injection of 3M MgCl2. Kinetic analysis of each reference subtracted injection series was performed using the BIAEvaluation software (Cytiva). All sensorgram series were fit to a 1:1 (Langmuir) binding model of interaction.

### RBD library generation and identification of escape mutants

Yeast display plasmids pJS697 and pJS699 used for surface display of Wuhan-Hu-1 S RBD N343Q were previously described^60^. Using these plasmids, 119 surface exposed positions on the original Wuhan-Hu-1 S RBD N343Q (positions 333-537) were mutated to every other amino acid plus stop codon using degenerate NNK primers using comprehensive nicking mutagenesis ^61^ exactly as previously described^36, 62^. Two tiles were constructed for compatibility with 250bp paired end Illumina sequencing (tile 1: positions 333-436; tile 2: positions 437-527). Libraries were transformed into *S. cerevisiae* EBY100 and stocks of 1e8 viable yeast cells in 1 mL were stored in yeast storage buffer (20 w/v % glycerol, 200 mM NaCl, 20 mM HEPES pH 7.5) at -80°C. Library coverage was confirmed by 250 bp paired end Illumina deep sequencing, with statistics reported in Supplementary Table 5.

S RBD escape mutants are identified by a competitive assay between a nAb and soluble ACE2 as fully described in Francino-Urdaniz et al^36^. Briefly, yeast cells are grown in SDCAA for 4 hr at 30°C, pelleted, and then induced in SGCAA for 22 hr at 22°C. Cells are washed thoroughly in PBSA (PBS containing 1g/L BSA) and then 3x10^7^ induced EBY100 yeast cells displaying S RBD were labelled with 10 !g/ml nAb IgG for 30min at room temperature with mixing by pipetting every 10min in PBSA.

The same cells were labelled with 75 nM chemically biotinylated ACE2, in the same tube, for 30min at room temperature in PBSA with mixing by pipetting every 10 min. The cells were centrifuged and washed with 1mL PBSA. Cells were then labeled with 1.2 μL FITC, 0.5 μL SAPE and 98.3 μL PBSA for 10min at 4°C. Cells were centrifuged, washed with 1mL PBSF, resuspended to 1 mL PBSA and sorted using FACS. Two additional populations were sorted: a reference population containing only an FSC/SSC gate for isolation of yeast cells (see below) and a control population of library not competitively labelled.

To discriminate cell populations FACS gates are used as shown in Figure S7: an FSC/SSC gate for isolation of yeast cells, FSC-H/FSC-A gate to discriminate single cells, a FSC-A/FITC+ gate selects the cells displaying the RBD on their surface and the top 2% of cells by a PE^+^/FITC^+^ gate is collected. At least 2.0x10^5^ cells are collected and recovered in SDCAA with 50 !g/mL Kanamycin and 1x Pen/Strep for 30h at 30°C. The collected DNA is sequenced using 250 bp PE on an Illumina MiSeq and analyzed with the code developed by Francino-Urdaniz et al^36^. Outputs from the code are a per-mutation enrichment ratio defined as the log2-transform of the change in frequency of the selected population relative to the reference population and a false discovery rate (FDR) as previously described^36^.

### Animal Study

Golden Syrian hamsters were provided by Charles River Laboratories (CRL:LVG(SYR)) and housed at the Scripps Research Institute. Animals were infused with antibodies intraperitoneally as described previously^23^. The Scripps Research Institutional Animal Care and Use Committee (IACUC) approved all experimental procedures involving all the animals in accordance with Protocol #20-0003.

### Viral load measurements - plaque assay

SARS-CoV-2 titers were measured by homogenizing lung tissue in DMEM 2% FCS using 100 µm cell strainers (Myriad, 2825-8367). Homogenized organs were titrated 1:10 over 6 steps and layered over Vero-E6 cells. After 1 h of incubation at 37°C, a 1% methylcellulose in DMEM overlay was added, and the cells were incubated for 3 days at 37°C. Cells were fixed with 4% PFA and plaques were counted by crystal violet staining.

### Statistical analysis

Statistical analysis was performed using Graph Pad Prism 8 for Mac, Graph Pad Software, San Diego, California, USA. Groups of data were compared using several methods including the grouped parametric One-Way ANOVA test and the grouped non-parametric Kruskall-Walli test. Data were considered statistically significant at p < 0.05.

## ACKNOWLEDGEMENTS

Research reported in this publication was supported by the National Institute of Allergy and Infectious Diseases of the National Institutes of Health under Award Number R01AI141452 to T.A.W. This work was supported by NIH CHAVD UM1 AI44462 (D.R.B.), the IAVI Neutralizing Antibody Center, the Bill and Melinda Gates Foundation OPP1170236 and INV-004923 (D.R.B., A.B.W.), and the James B. Pendleton Charitable Trust (D.R.B.).

## AUTHOR CONTRIBUTIONS

F.Z. and J.G.J. conceived and designed the study. F.Z. and S.B. performed yeast library preparation, yeast cell staining, sorting, sequencing, and cloning experiments. F.Z., S.B., A.B., and O.L. expressed and purified the monoclonal antibodies. G.O., and H.T. performed Cryo-EM sample preparation, imaging, data processing and model building. F.Z., S.B., and A.B. characterized monoclonal antibodies in functional assays and biophysical analysis. A.B. and O.L. generated mutant RBD protein plasmids and expressed recombinant RBD and S proteins. J.W. performed surface plasmon resonance experiment and analysis. A.B., S.B., P.Z. and L.P. generated pseudovirus mutant constructs, produced pseudovirus and performed neutralization assay. I.F.U and T.A.W. generated RBD mutant library, analyzed and identified escaped mutants. C.K. and N.S. performed hamster protection experiments including antibody infusion, virus challenge, weight analysis, viral load measurement. F.Z., C.K., G.O., D.R.B., and J.G.J. wrote the manuscript and all authors reviewed and edited the manuscript.

---

**Supplementary Fig. 1.**
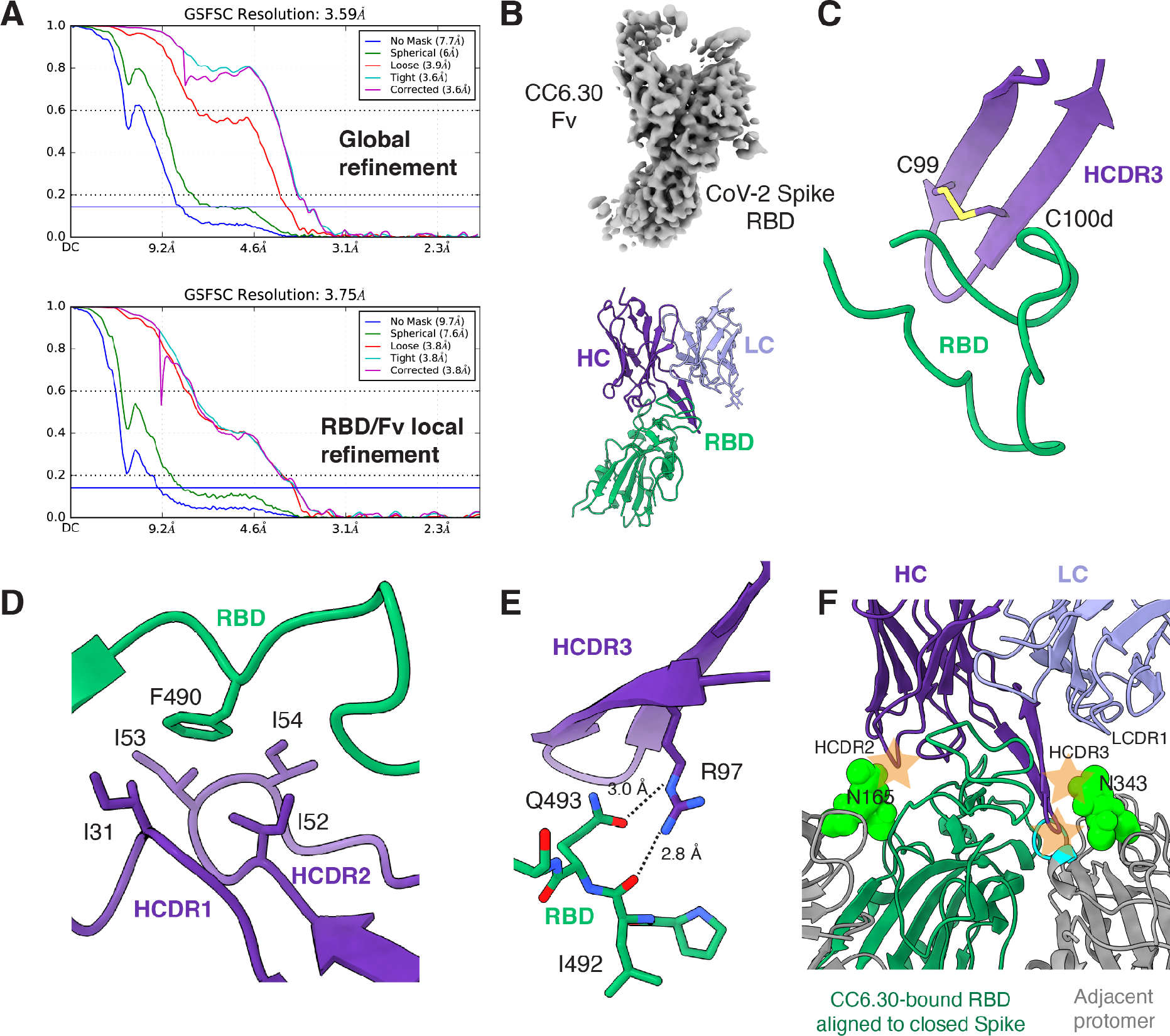
CryoEM reconstruction of nAb CC6.30 in complex with SARS-CoV-2 HPM7 Spike. Related to Fig.1**. a**, Fourier shell correlation (FSC) of global refinement (*top*) and RBD/Fv local refinement (*bottom*) of the CC6.30:HPM7 complex. **b**, Local refinement map (*top*) and model (*bottom*). **c**, CC6.30 HCDR3 intrachain disulfide bond. **d**, Hydrophobic interactions between RBD F490 and multiple CC6.30 heavy chain residues. **e**, Putative hydrogen bonds (dashed lines) between CC6.30 HCDR3 R97 and RBD. **f**, Superposition of RBD:CC6.30 onto a closed (3 down RBDs) Spike predicts multiple clashes (orange stars) with glycans (N165, N343) and peptide (residues 367-371; cyan) with the adjacent protomer.

**Supplementary Fig. 2.**
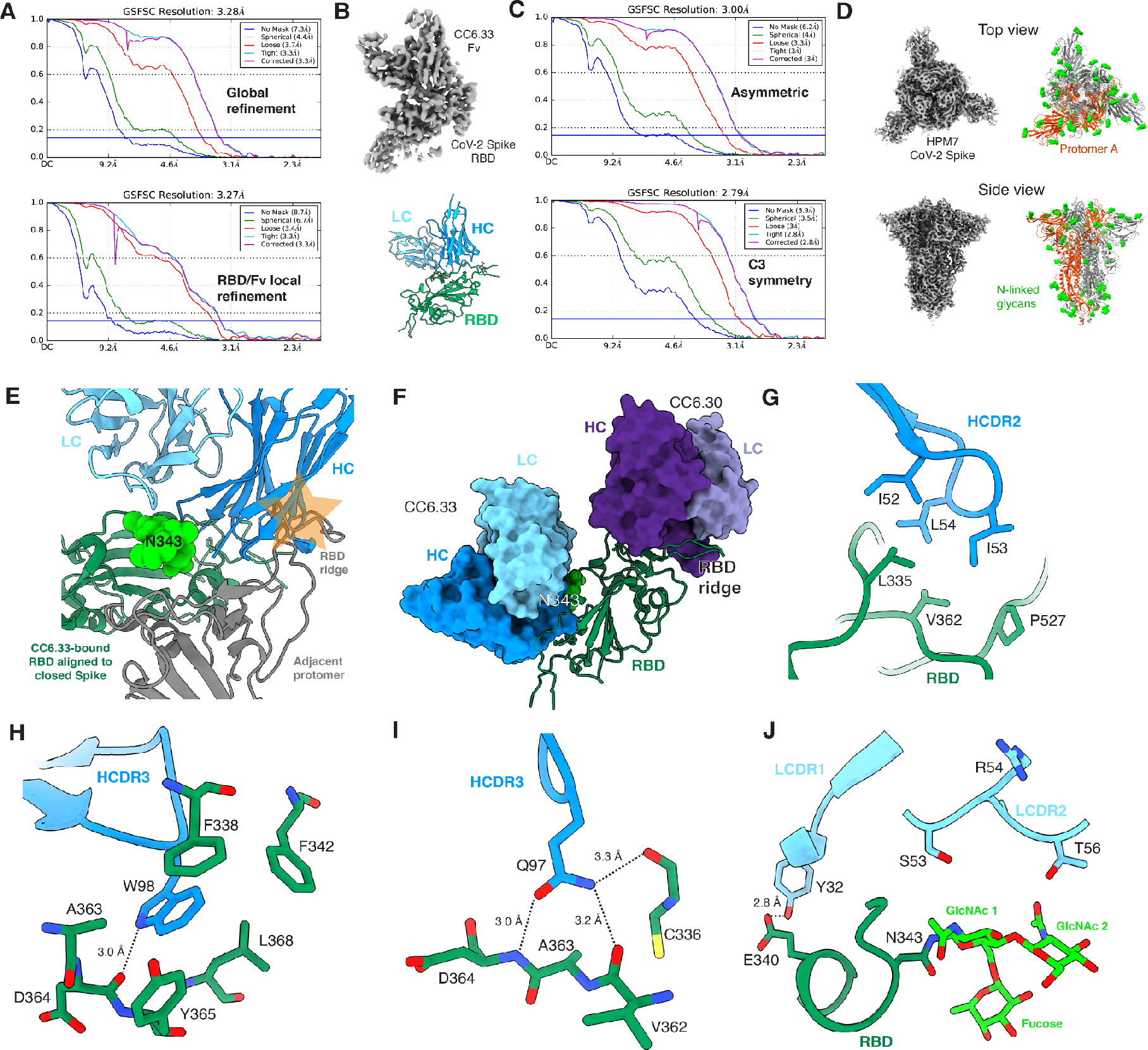
CryoEM reconstruction of nAb CC6.33 in complex with SARS-CoV-2 HPM7 Spike. Related to Fig.2**. a**, Fourier shell correlation (FSC) of global refinement (*top*) and RBD/Fv local refinement (*bottom*) of the CC6.33:HPM7 complex. **b**, Local refinement map (*top*) and model (*bottom*). **c**, Fourier shell correlation (FSC) of asymmetric (*top*) and C3-symmetric (*bottom*) ligand-free HPM7 SARS-CoV-2 Spike reconstructions. **d**, Map and model of ligand-free HPM7 SARS-CoV-2 Spike C3-symmetric reconstruction. **e**, Superposition of RBD:CC6.33 onto a closed (3 down RBDs) Spike predicts a clash with the RBD ridge of an adjacent protomer that is relieved with slight opening of the apex. **f**, Modeling of CC6.30 and CC6.33 on the same RBD reveals non-overlapping epitopes. **g**, Hydrophobic packing between CC6.33 HCDR2 and the RBD. **h**, Interactions between CC6.33 HCDR3 Trp98 and the RBD. **i**, Putative hydrogen bonding between CC6.33 HCDR3 Gln97 and the RBD. **j**, Interface between CC6.33 LCDR1 and LCDR2, and RBD, including glycan N343.

**Supplementary Fig. 3.**
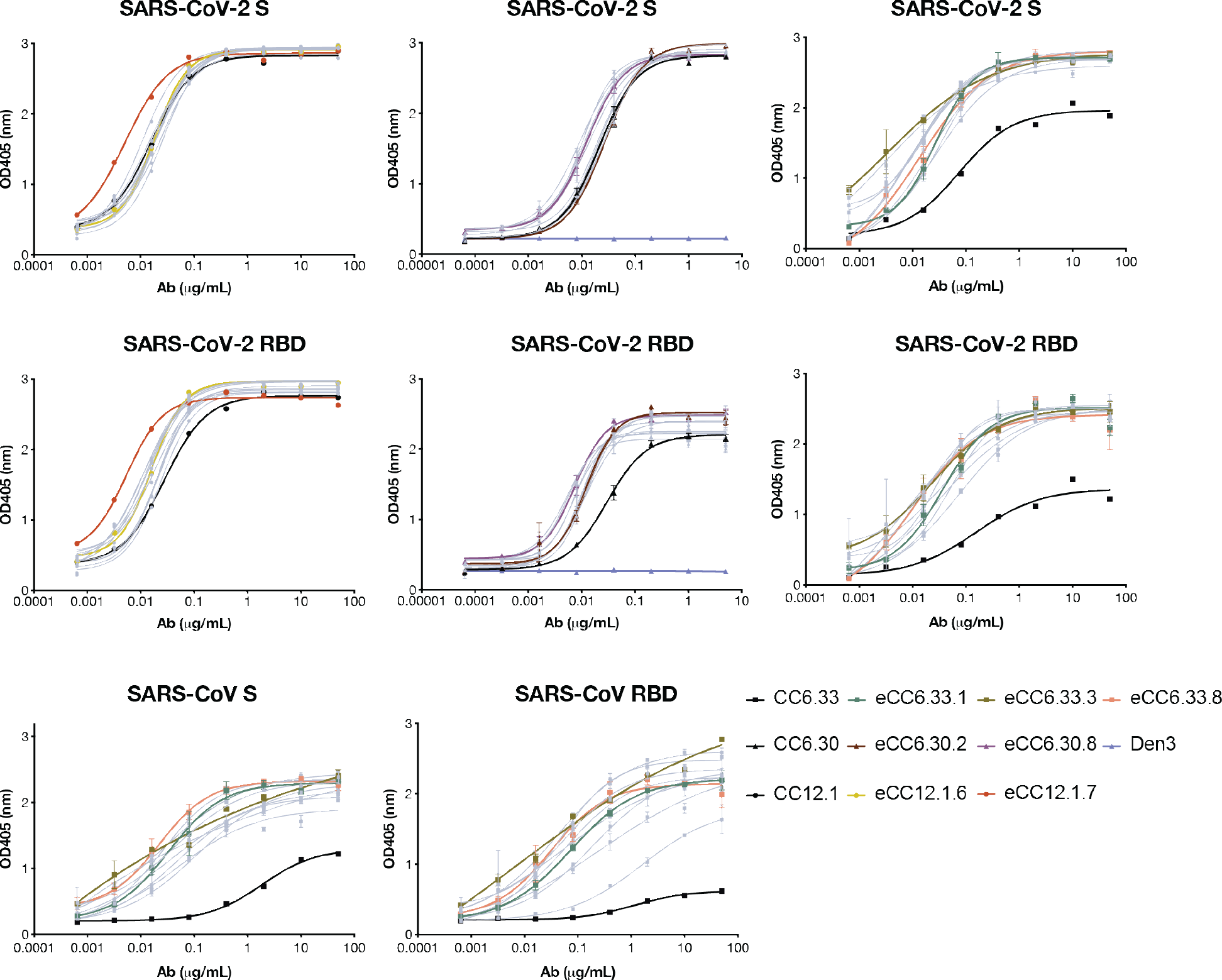
Engineered antibody variants ELISA binding to SARS-CoV-2 and SARS-CoV-1 S/RBD proteins. Related to Fig.3. Parental and engineered Abs were evaluated binding against his-tagged recombinant S/RBD protein. Parental mAbs were colored in black while some key enAbs were colored according to the key. The rest of enAbs were colored in grey. Antibodies were tested in duplicates. Error bars represent standard deviations.

**Supplementary Fig. 4.**
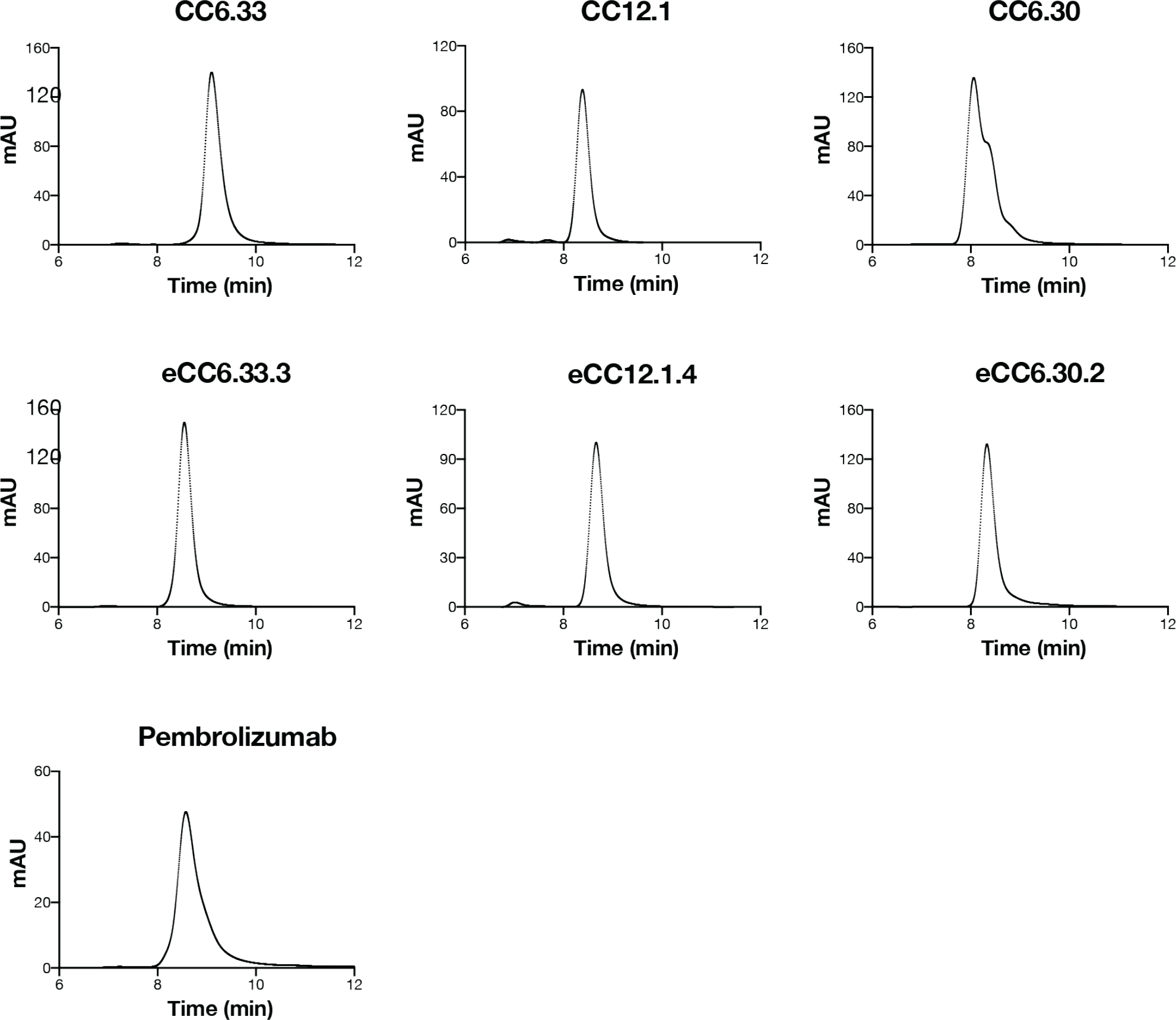
Size exclusion chromatography analysis of parental and engineered SARS-CoV-2 mAbs. Related to Fig.3. The antibodies were analyzed by size exclusion chromatography using the 1260 Infinity II (Agilent). 15 uL of each antibody was injected into the TSKgel SuperSW mAb column at 2 mg/mL. An FDA-approved mAb Pembrolizumab was included as control.

**Supplementary Fig. 5.**
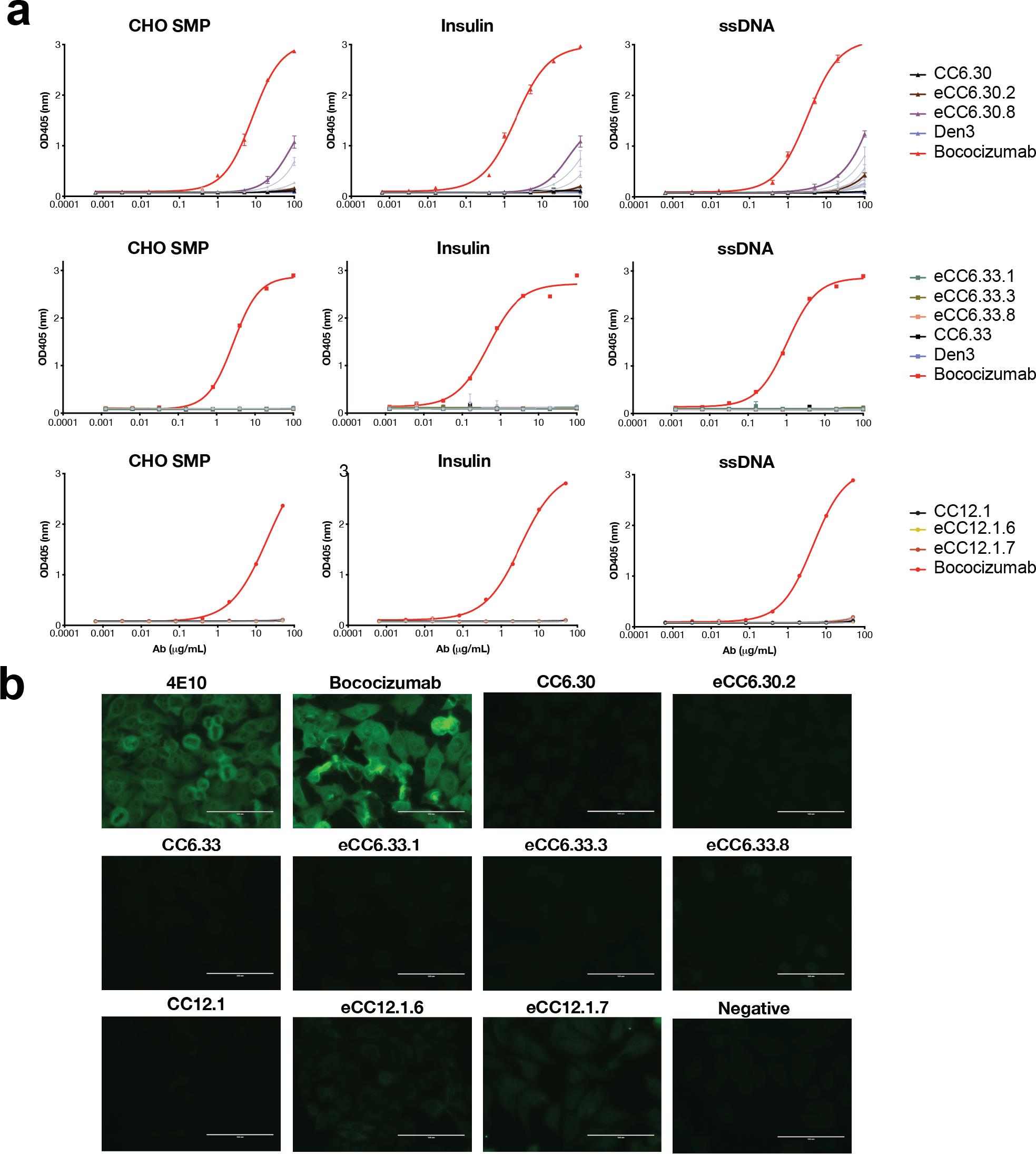
Polyreactivity of parental and enhanced nAbs by SPR. Related to Fig.3**. a**, ELISA of eCC6.30, eCC6.33, eCC12.1 variants and parental clones to CHO solubilized membrane proteins, human insulin, and ssDNA. Bococizumab serves as positive control while Den3 serves as negative control. Error bars represent standard deviations. **b**, HEp2 epithelial cells staining with mAbs at 100 ug/mL. 4E10 and Bococizumab serve as positive control.

**Supplementary Fig. 6.**
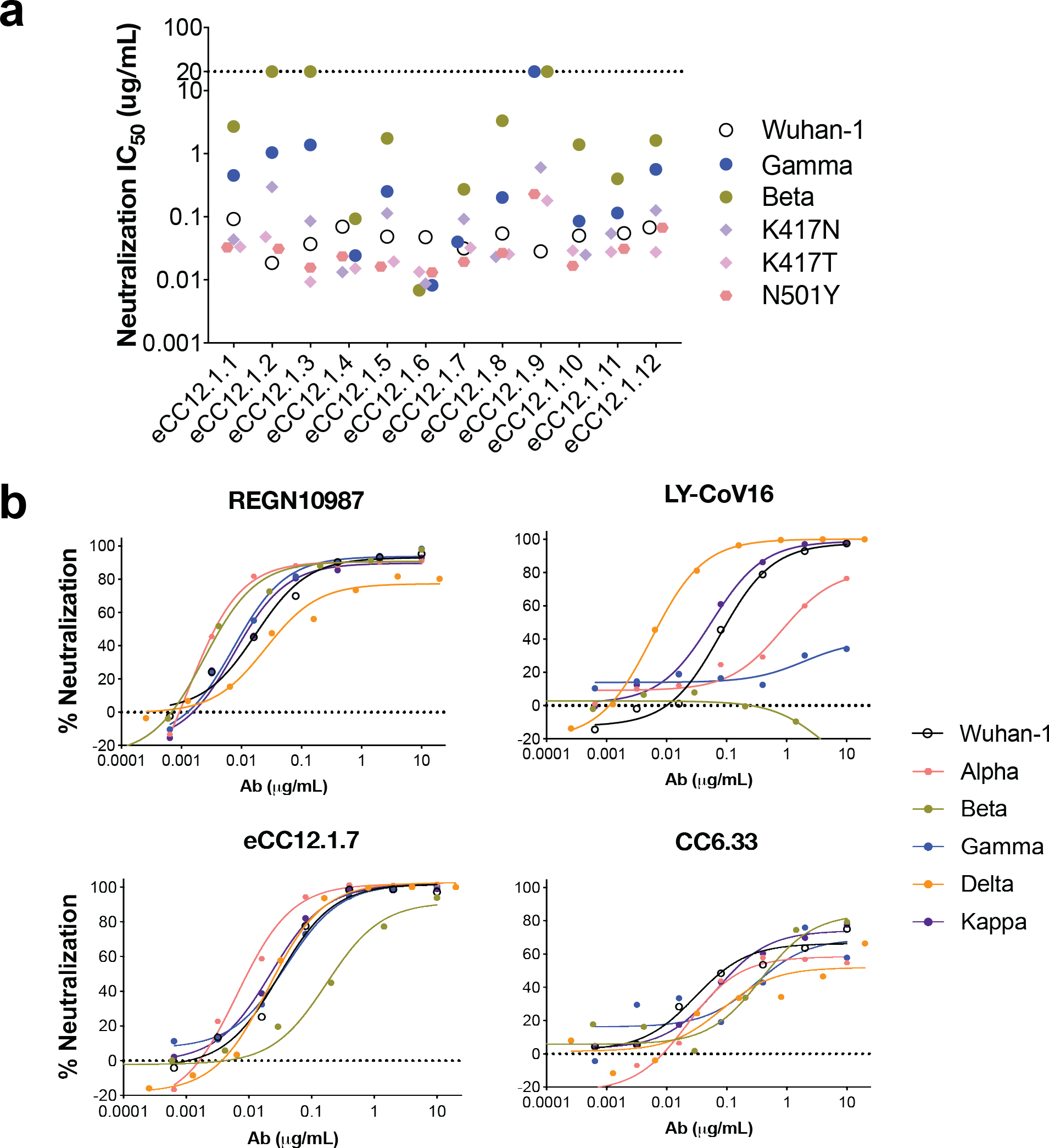
Neutralization of mAbs against VOCs. Related to Fig.4**. a**, Neutralization IC50 of 12 eCC12.1 variants against Wuhan-1, K417N, K417T, N501Y, Beta, and Gamma variants. Antibody started at 20 ug/mL and was tested in duplicates. **b**, Neutralization curves of REGN10987, LY-CoV16, CC6.33 and eCC12.1.7 against Wuhan-1 as well as VOCs. Virus strains were colored according to the key.

**Supplementary Fig. 7.**
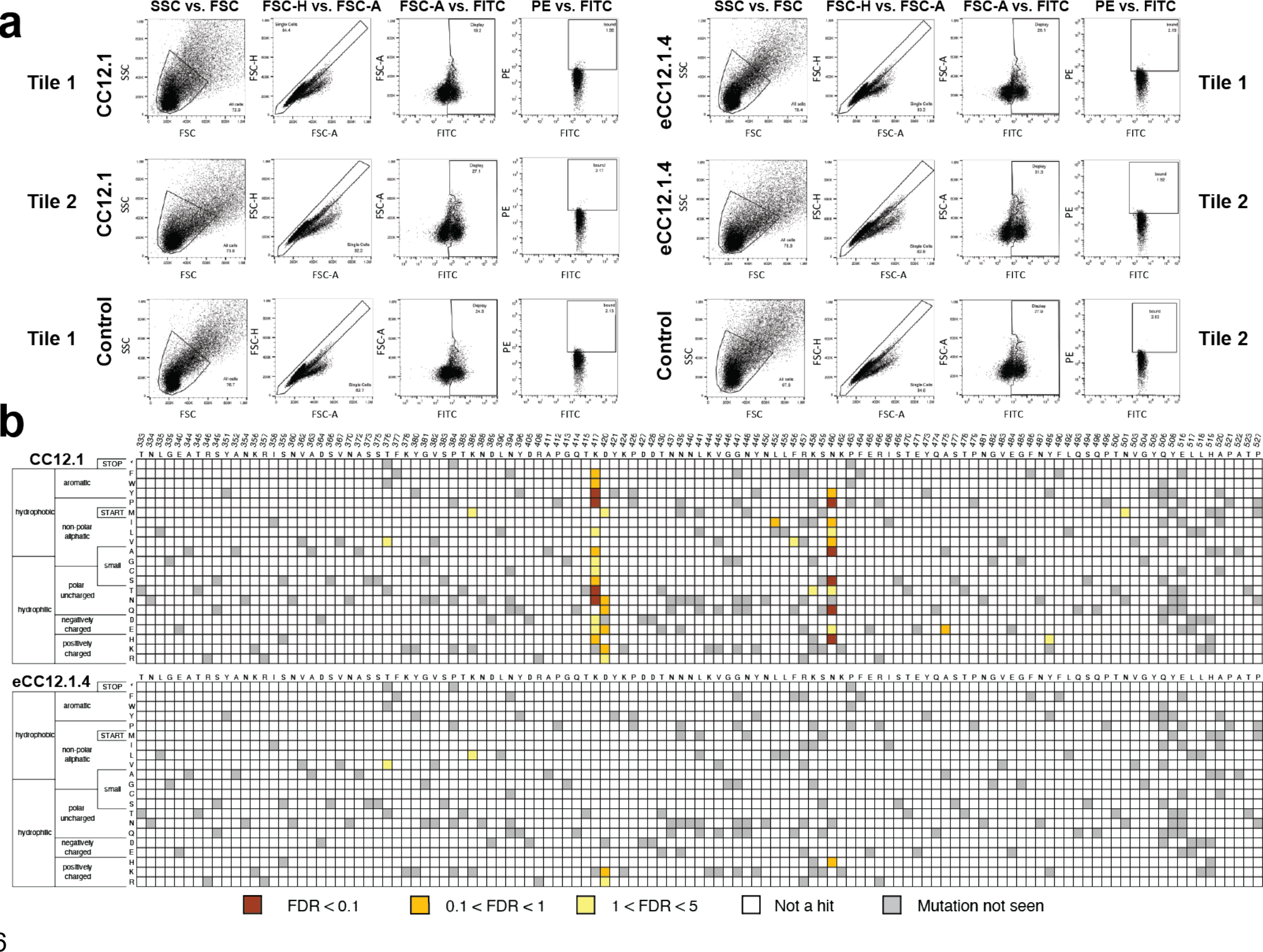
Identification of SARS-CoV-2 RBD escape mutants using yeast screening. Related to Fig.5**. a**, Gating strategy and gates set for the escape mutant analysis for CC12.1 and eCC12.1.4. Control is the population of yeast cells without labeling by either biotinylated ACE2 or a given nAb. **b**, Heatmap showing predicted RBD (residues 333-527) escape mutants for CC12.1 and eCC12.1.4 in yellow to burnt orange with varying levels of confidence according to the key.

**Supplementary Fig. 8.**
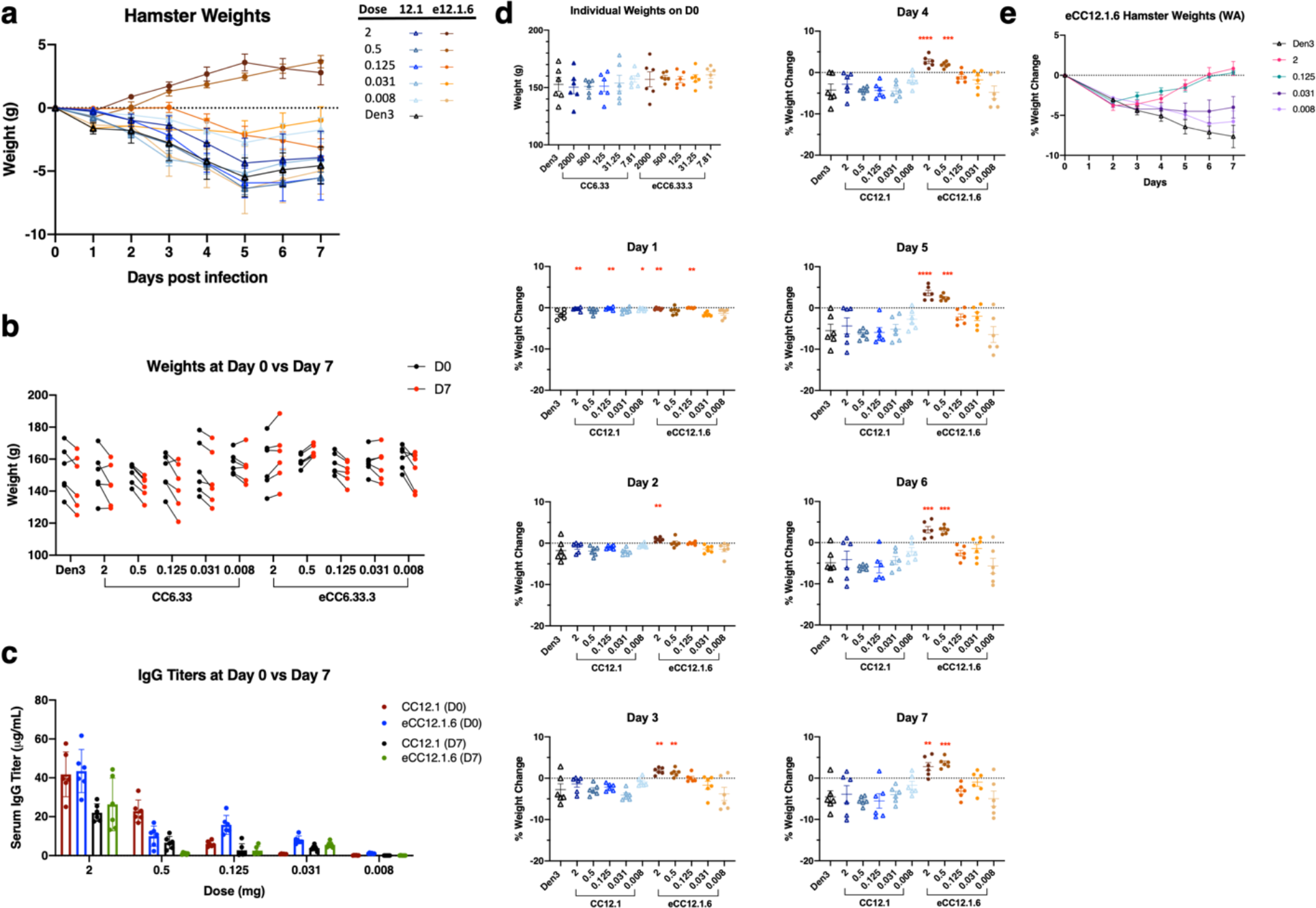
Supplemental Animal Protection Studies, CC12.1 and eCC12.1.6. Related to Fig.6**. a**, Weight trends of all groups included in the CC12.1 vs eCC12.1.6 prophylactic protection study. **b**, Weights of animals at time of challenge (Day 0) compared to weights at time of sacrifice (Day 7). **c**, Serum human IgG concentration at time of infection (Day 0) compared to sacrifice (Day 7). **d**, Percent weight loss by day compared to weights recorded at time of infection at day 0. **e**, Weight trend of animals administered with eCC12.1.6 and subsequently challenged with 1 × 10^5^ PFU of SARS-CoV-2 (USA-WA1/2020) three days later.

**Supplementary Fig. 9.**
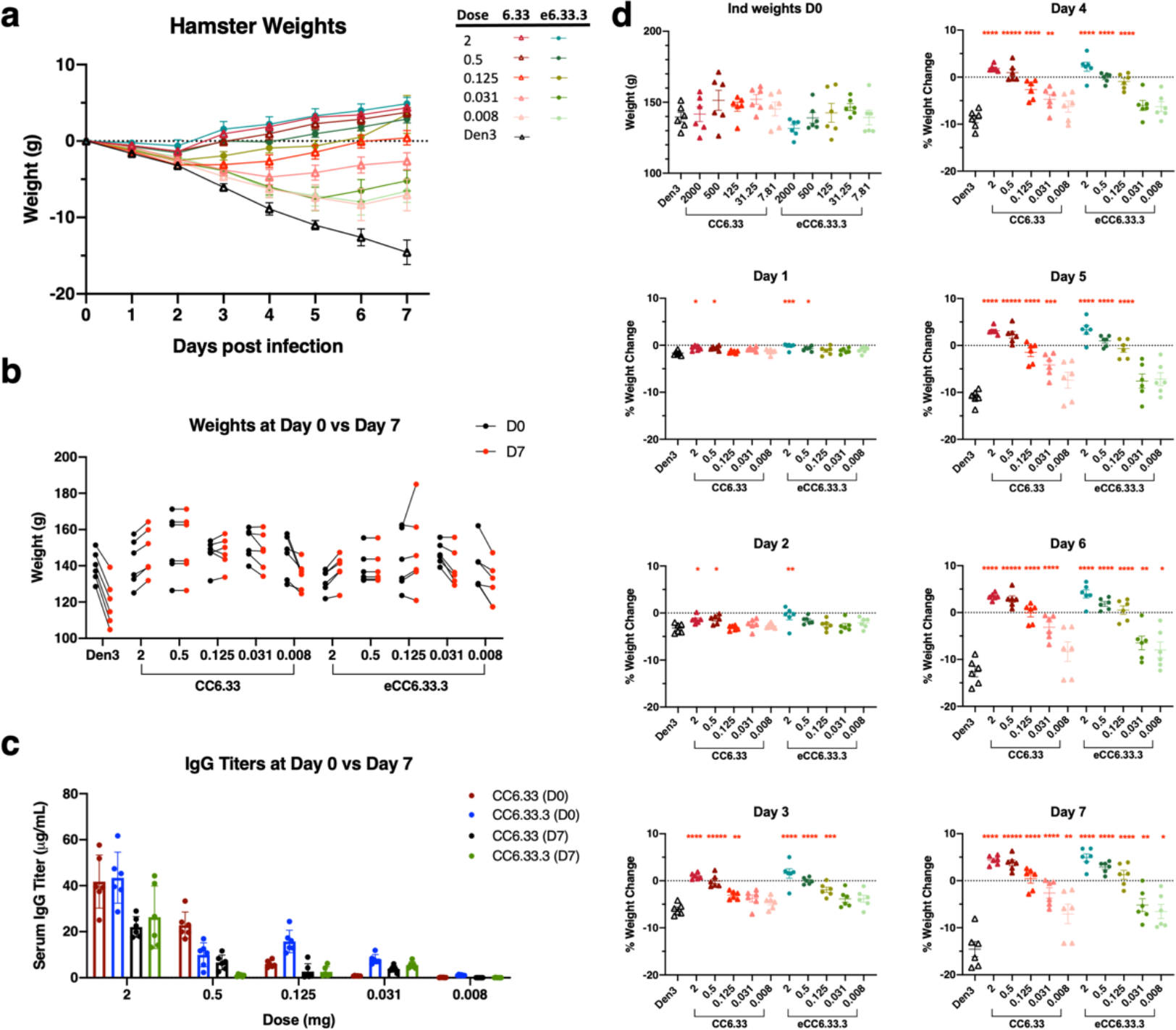
Supplemental Animal Protection Studies, CC6.33 and eCC6.33.3. Related to Fig.6**. a**, Weight trends of all groups included in the CC6.33 vs eCC6.33.3 protection study. **b**, Weights of animals at time of challenge (Day 0) compared to weights at time of sacrifice (Day 7). **c**, Serum human IgG concentration at time of infection (Day 0) compared to sacrifice (Day 7). **d**, Percent weight loss by day compared to individual weights recorded at time of infection at day 0.

**Supplementary Fig. 10.**
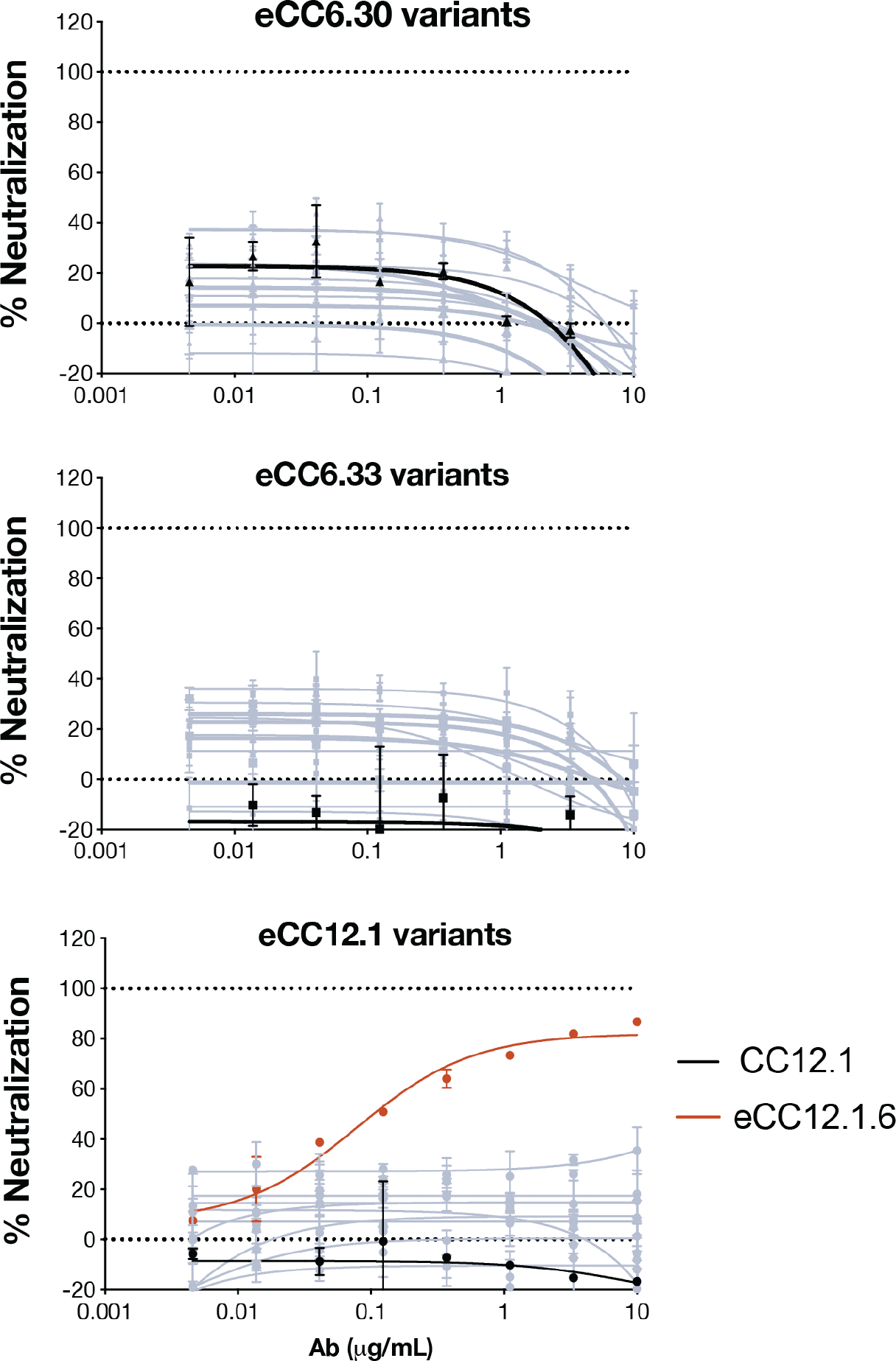
Neutralization of enAbs against the Omicron variant. Related to Fig.7. Neutralization curves of eCC6.33, eCC6.30, and eCC12.1 variants against pseudotyped Omicron VOC. Parental antibodies were highlighted in black whereas enAbs were in grey. eCC12.1.6 was highlighted according to the key. Error bars represent standard deviations

**Supplementary Table 1.** CryoEM data collection, refinement and model building statistics (CC6.30 dataset). Related to Fig.1.

| Map | CC6.30 Fab +<br>SARS-CoV-2 S<br>HPM7 (Non-uniform refinement) | CC6.30 Fab +<br>SARS-CoV-2 S<br>HPM7 (RBD/Fv<br>local refinement) |
| --- | --- | --- |
| EMDB | EMD-24697 | EMD-24699 |
| **Data collection** |  |  |
| Microscope | FEI Titan Krios | FEI Titan Krios |
| Voltage (kV) | 300 | 300 |
| Detector | Gatan K2 Summit | Gatan K2 Summit |
| Recording mode | Counting | Counting |
| Nominal magnification | 29,000 | 29,000 |
| Movie micrograph pixel size (Å) | 1.03 | 1.03 |
| Dose rate (e<sup>-</sup>/[(camera pixel)*s]) | 5.414 | 5.414 |
| Number of frames per movie micrograph | 39 | 39 |
| Frame exposure time (ms) | 250 | 250 |
| Movie micrograph exposure time (s) | 9.75 | 9.75 |
| Total dose (e<sup>-</sup>/Å<sup>2</sup>) | 50 | 50 |
| Defocus range (µm) | -0.6 to -1.5 | -0.6 to -1.5 |
| **EM data processing** |  |  |
| Number of movie micrographs | 3,348 | 3,348 |
| Number of molecular projection images in map | 55,248 | 94,532 |
| Symmetry | C1 | C1 |
| Map resolution (FSC 0.143; Å) | 3.6 | 3.8 |
| Map sharpening B-factor (Å<sup>2</sup>) | -79.9 | -50 |
| **Structure Building and Validation** |  |  |
| *Number of atoms in deposited model* |  |  |
| SARS-CoV-2 S protein | 24,577 | 1,388 |
| Glycans | 560 | 42 |
| CC6.30 Fv | 3,536 | 1,768 |
| MolProbity score | 0.91 | 1.19 |
| Clashscore | 0.81 | 0.96 |
| Map correlation coefficient | 0.72 | 0.72 |
| EMRinger score | 2.57 | 2.53 |
| *RMSD from ideal* |  |  |
| Bond length (Å) | 0.02 | 0.02 |
| Bond angles (°) | 1.81 | 1.88 |
| *Ramachandran plot* |  |  |
| Favored (%) | 97.12 | 93.92 |
| Allowed (%) | 2.88 | 6.08 |
| Outliers (%) | 0 | 0 |
| Side chain rotamer outliers (%) | 0.06 | 0 |
| PDB | 7ru5 | 7ru8 |

**Supplementary Table 2.** CryoEM data collection, refinement and model building statistics (CC6.33 dataset). Related to Fig.2.

| Map | CC6.33 IgG + SARS-CoV-2 S HPM7 (Non-uniform refinement) | CC6.33 IgG + SARS-CoV-2 S HPM7 (RBD/Fv local refinement) | SARS-CoV-2 S HPM7 (C3 symmetry) | SARS-CoV-2 S HPM7 (C1 symmetry) |
| --- | --- | --- | --- | --- |
| EMDB | EMD-24695 | EMD-24696 | EMD-24693 | EMD-24694 |
| **Data collection** |  |  |  |  |
| Microscope | FEI Titan Krios | FEI Titan Krios | FEI Titan Krios | FEI Titan Krios |
| Voltage (kV) | 300 | 300 | 300 | 300 |
| Detector | Gatan K2 Summit | Gatan K2 Summit | Gatan K2 Summit | Gatan K2 Summit |
| Recording mode | Counting | Counting | Counting | Counting |
| Nominal magnification | 29,000 | 29,000 | 29,000 | 29,000 |
| Movie micrograph pixel size (Å) | 1.03 | 1.03 | 1.03 | 1.03 |
| Dose rate (e<sup>-</sup>/[(camera pixel)*s]) | 5.414 | 5.414 | 5.414 | 5.414 |
| Number of frames per movie micrograph | 39 | 39 | 39 | 39 |
| Frame exposure time (ms) | 250 | 250 | 250 | 250 |
| Movie micrograph exposure time (s) | 9.75 | 9.75 | 9.75 | 9.75 |
| Total dose (e<sup>-</sup>/Å<sup>2</sup>) | 50 | 50 | 50 | 50 |
| Defocus range (μm) | -0.5 to -1.5 | -0.5 to -1.5 | -0.5 to -1.5 | -0.5 to -1.5 |
| **EM data processing** |  |  |  |  |
| Number of movie micrographs | 3,465 | 3,465 | 3,465 | 3,465 |
| Number of molecular projection images in map | 47,461 | 96,012 | 55,575 | 55,575 |
| Symmetry | C1 | C1 | C3 | C1 |
| Map resolution (FSC 0.143; Å) | 3.3 | 3.3 | 2.8 | 3 |
| Map sharpening B-factor (Å<sup>2</sup>) | -59.4 | -33.9 | -66.9 | -54.6 |
| **Structure Building and Validation** |  |  |  |  |
| *Number of atoms in deposited model* |  |  |  |  |
| SARS-CoV-2 S protein | 24,175 | 1,610 | 25,737 | 25,778 |
| Glycans | 584 | 52 | 756 | 868 |
| CC6.33 Fv | 3,398 | 1,720 | 0 | 0 |
| MolProbity score | 1 | 0.89 | 0.91 | 0.83 |
| Clashscore | 1.04 | 0.6 | 0.77 | 0.46 |
| Map correlation coefficient | 0.81 | 0.8 | 0.83 | 0.83 |
| EMRinger score | 2.55 | 3.98 | 5.05 | 4.15 |
| *RMSD from ideal* |  |  |  |  |
| Bond length (Å) | 0.02 | 0.02 | 0.02 | 0.02 |
| Bond angles (°) | 1.85 | 1.9 | 1.77 | 1.78 |
| *Ramachandran plot* |  |  |  |  |
| Favored (%) | 96.82 | 96.9 | 96.97 | 97.04 |
| Allowed (%) | 3.18 | 3.1 | 3.03 | 2.96 |
| Outliers (%) | 0 | 0 | 0 | 0 |
| Side chain rotamer outliers (%) | 0.32 | 0.55 | 0 | 0.14 |
| PDB | 7ru3 | 7ru4 | 7ru1 | 7ru2 |

**Supplementary Table 3.**
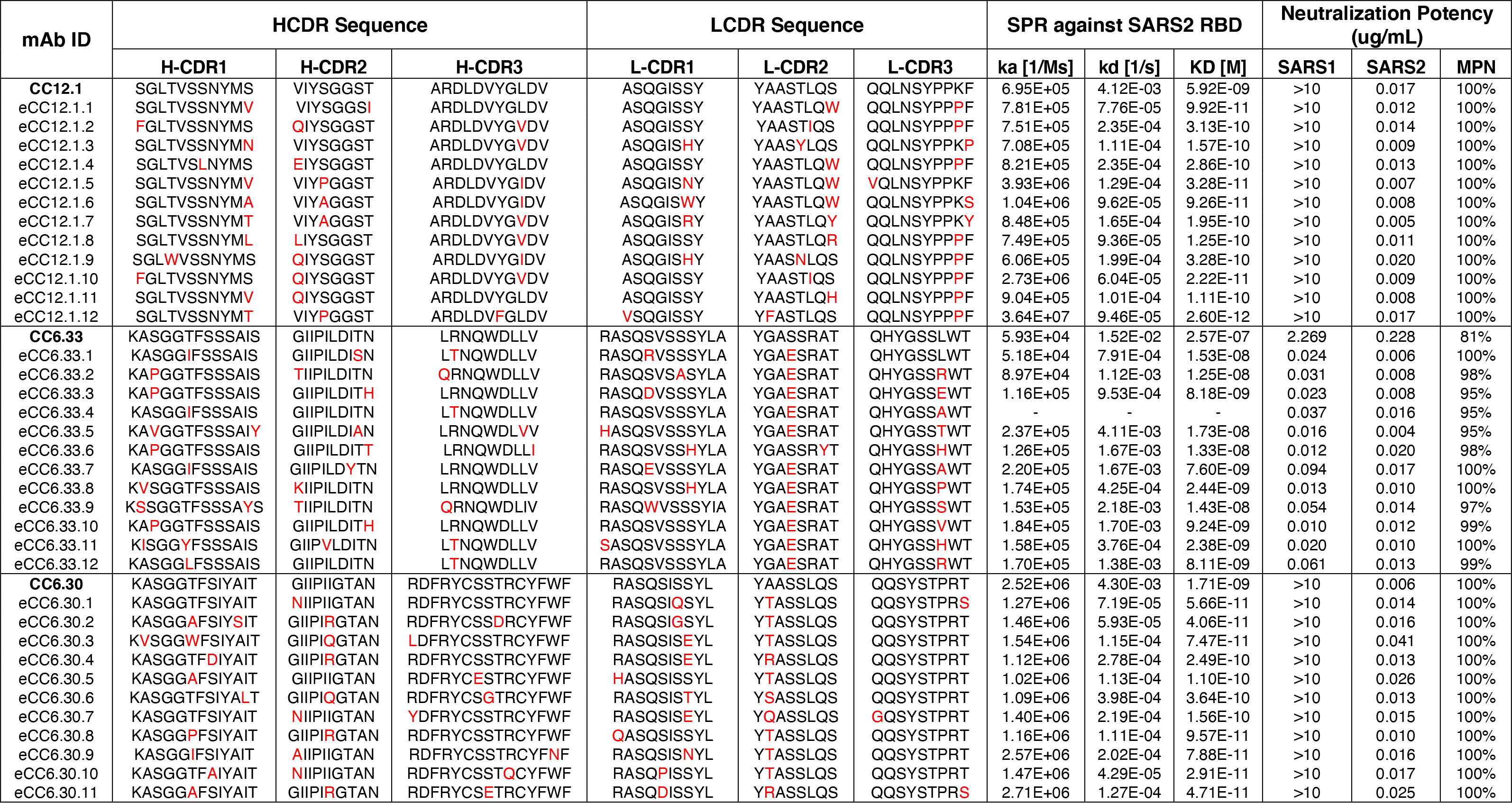
CDR loop sequences, RBD binding affinity, and neutralization potency of parental and engineered antibodies. Related to Fig.3. Summary table of parental CC12.1, CC6.33, CC6.30 and 35 engineered antibodies with sequences of 6 CDR loops, binding affinity against SARS-CoV-2 RBD, SARS-CoV-1/2 neutralization potency and maximum percentage of neutralization (MPN). Mutations of engineered antibodies at CDR loops were highlighted in red. Antibodies were captured via Fc-capture to an anti-human IgG Fc antibody and varying concentrations of SARS-CoV-2 RBD were injected using a multi-cycle method. Association and dissociation rate constants calculated through a 1:1 Langmuir binding model using the BIAevaluation software. Neutralization assay was performed using pseudotyped SARS-CoV-1 and SARS-CoV-2 with Hela-hACE2 cell line. All antibodies were tested in duplicates.

**Supplementary Table 4.**
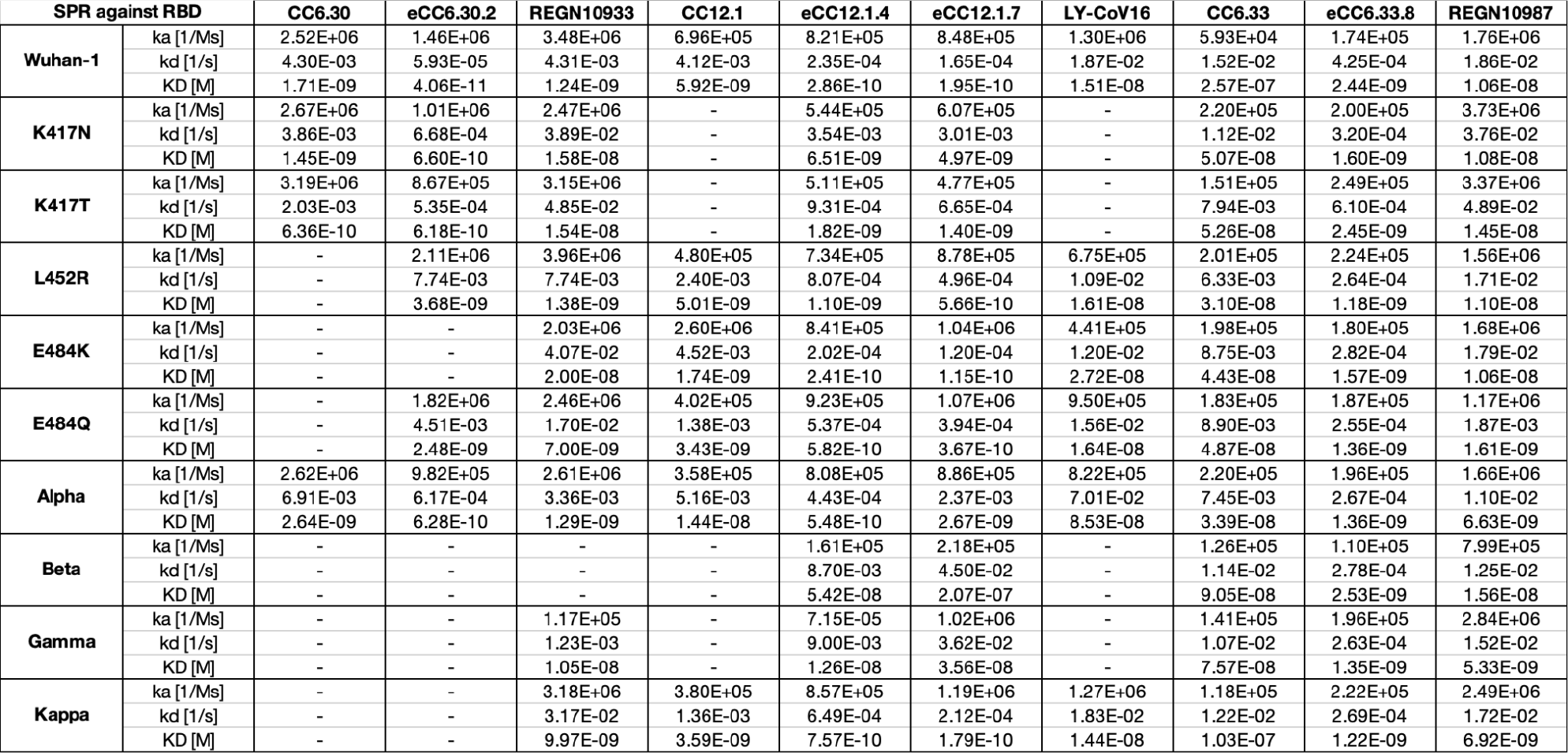
Surface plasmon resonance of SARS-CoV-2 nAbs against RBD variants. Related to Fig.4. Antibodies were captured via Fc-capture to an anti-human IgG Fc antibody and varying concentrations of Wuhan-1 RBD or SARS-CoV-2 RBD variants were injected using a multi-cycle method. Association and dissociation rate constants calculated through a 1:1 Langmuir binding model using the BIAevaluation software. Antibodies that did not bind to RBD variants or their binding curves did not fit into the model were shown as no values in the table.

**Supplementary Table 5.** S RBD Single saturation mutagenesis library coverage statistics. Related to Fig.5.

| Tile Number | Tile 1 | Tile 2 |
| --- | --- | --- |
| Positions | 333-436 | 437-537 |
| Number of Designed Mutations | 1120 | 1260 |
| *E. coli* transformants Obtained from<br>Nicking Saturation Mutagenesis | 5.0E+05 | 1.5E+06 |
| Yeast transformants Obtained from<br>Homologous Recombination | 8.0E+05 | 1.0E+06 |
| Library Coverage Per Tile | 97%<br>(1089/1120) | 92%<br>(1161/1260) |
| Overall Library Coverage | 94.5% (2250/2380) |  |

